# What drives variation in conspecific negative density-dependence? A demographic perspective

**DOI:** 10.64898/2026.07.21.739908

**Authors:** Daniel J. B. Smith, Dale Forrister, Brian E. Sedio, Annette Ostling

## Abstract

Conspecific negative density dependence (CNDD), the reduced survival of juvenile trees at high conspecific density, is considered a key force maintaining plant diversity. Variation in CNDD across species and environments is typically attributed to variation in sensitivities to natural enemies (e.g., pathogen transmission/virulence) or sensitivity to intraspecific competition. We show that the density-independent component of vital rates — baseline mortality and growth — substantially alters the cumulative consequences of conspecific density on survival, independently of any change in instantaneous sensitivity to conspecific crowding. This reflects genuine demographic effects on opportunities for crowding to shape survival, and on population characteristics shaping the negative influence of conspecifics, such as infection prevalence and size structure. It is not an artifact of how CNDD is measured. Our results, derived from a pathogen model and a size-structured seedling model, provide a demographic framework for assessing potential drivers of CNDD variation among species and across abiotic gradients.

## Introduction

Tree communities display striking variation in diversity; understanding the mechanisms that maintain this variation is a central goal in ecology. One prominent idea is that juvenile trees often survive poorly at high local conspecific density. This pattern, commonly termed Conspecific Negative Density Dependence (CNDD), is often attributed to specialized natural enemies such as pests and pathogens that accumulate in areas of high host abundance (Janzen, 1970; Connell, 1971; Comita et al., 2014; Bever et al., 2015). If juveniles of common species experience lower survival than those of rarer species, CNDD can generate a rare-species advantage and contribute to species coexistence (Chesson, 2000). Variation in CNDD — both between species and on abiotic gradients — is thought to regulate how CNDD maintains diversity (LaManna et al., 2024). However, the processes shaping variation in CNDD strength remain poorly understood.

CNDD that influences coexistence is best understood not as an instantaneous per-capita effect of neighbors on a focal individual, but as the cumulative survival consequences that conspecific crowding generates over ecologically meaningful intervals of juvenile life (Fig. 1A-D,F). Specifically, recruitment from seed to adulthood can be understood as the compound probability of surviving each of a series of successive intervals, each representing a distinct life-history stage (Fig. 1F). Over these intervals, juvenile trees experience time-varying hazards (Zens and Peart, 2003). In particular, as neighbors die or recruit, conspecific density changes and with it the per-capita hazard experienced by focal individuals; individuals also grow through size classes that carry different mortality risks. It is within this demographic context that CNDD is realized (Fig. 1B-D). Crucially, as we will argue, density-independent juvenile mortality and growth alter density-dependent processes. For example, if pathogen transmission depends on local seedling density, then baseline (density-independent) seedling mortality will change transmission dynamics; similarly, if density-dependent effects depend on neighbor size, then baseline growth will modify the strength of those effects (Fig. 1D, “example”). In this way, baseline demography can shape the cumulative survival consequences of conspecific crowding — what we term *realized interval CNDD* (Fig. 1D).

**Figure 1.**
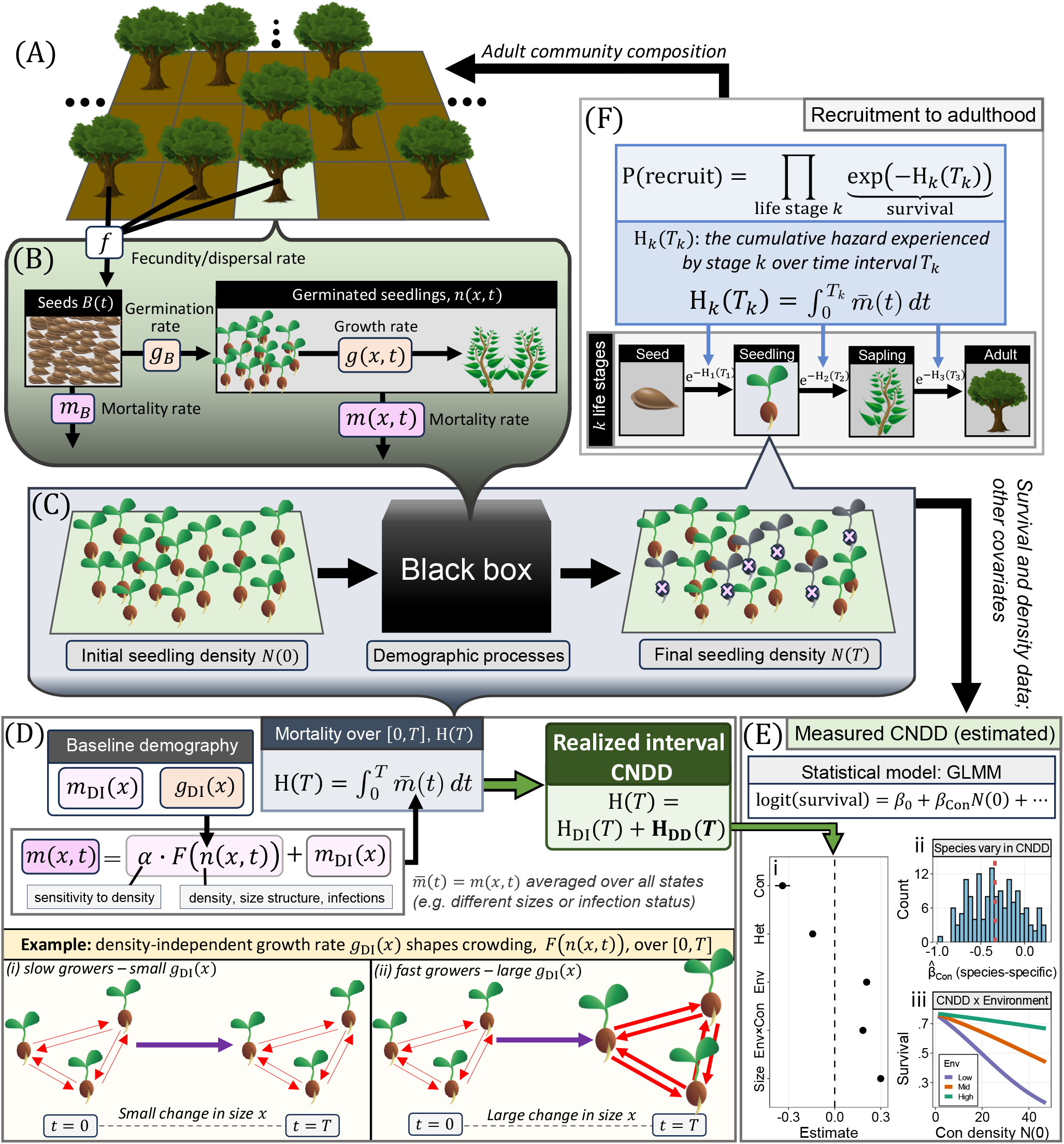
CNDD is an integrated effect of demographic processes. (A) Schematic landscape with a focal location (green). (B) Key demographic processes: adults contribute seeds at rate *f* ; seeds *B*(*t*) die at rate *m*_*B*_ and germinate at rate *g*_*B*_ to produce a size-structured seedling cohort *n*(*x, t*). Seedlings grow at rate *g*(*x, t*) and die at rate *m*(*x, t*), both depending on size *x* and local density. (C) Observations over a census interval: seedling density is *N* (0) at time *t* = 0; after interval *T*, survival outcomes are recorded. The demo-graphic processes in (B) constitute a “black box” through which conspecific crowding effects accumulate. (D) The pipeline generating realized interval CNDD. Baseline mortality and growth rates (*m*_DI_(*x*), *g*_DI_(*x*)) shape how the cohort *n*(*x, t*) evolves over [0, *T* ], entering the instantaneous mortality rate *m*(*x, t*) = *α* · *F* (*n*(*x, t*)) + *m*_DI_(*x*). *m*(*x, t*) contains a density-dependent component governed by instantaneous density sensitivity (*α*) and a function of density *F* (*n*(*x, t*)). Averaging *m*(*x, t*) over the cohort’s size distribution gives 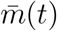, whose integral over [0, *T* ] is the cumulative hazard H(*T* ). Realized interval CNDD is the excess mortality attributable to density dependence, H_DD_(*T* ) = H(*T* ) − H_DI_(*T* ) (see Eq. 13). **Example:** assuming density-dependent effects increase with size, baseline growth rate *g*_DI_(*x*) shapes crowding *F* (*n*(*x, t*)) over [0, *T* ]: starting from identical cohorts at *t* = 0, slow growers (small *g*_DI_(*x*); i) change little in size *x* over *T*, whereas fast growers (large *g*_DI_(*x*); ii) grow substantially, increasing crowding. Arrow thickness represents crowding strength. (E) Measured CNDD, as commonly quantified with a GLMM. (i) Fixed effects describe survival as a function of conspecific density, heterospecific density, environment, their interactions, and size. (ii) Interspecific variation in measured CNDD summarized by species-specific random slopes 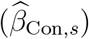. (iii) Predicted survival–density relationships across environments. (F) Recruitment to adulthood as the product of survival probabilities across life stages. Processes in (B–D) influence survival outcomes.

Current theory does not directly examine how CNDD emerges from the demographic processes unfolding within juvenile life. Most existing theoretical CNDD-related work examines how CNDD influences species coexistence (e.g., Adler and Muller-Landau, 2005; Chisholm and Muller-Landau, 2011; Sedio and Ostling, 2013; Stump and Chesson, 2015; Chisholm and Fung, 2020; May et al., 2020; Smith, 2022*a*). In these models, CNDD is represented by a single phenomenological parameter linking juvenile performance in recruiting to adulthood to conspecific density or distance — a fixed scalar that contains no demographic information. While this abstraction is instrumental for understanding how CNDD maintains coexistence, it prohibits any examination of how demographic processes modulate CNDD strength. Yet we argue that CNDD is shaped by exactly these demographic processes, through which conspecific effects accumulate over the time interval of the potential recruitment event (Fig. 1F).

The way CNDD is empirically measured similarly obscures these demographic processes. CNDD is measured from censuses in which neighborhood conditions are recorded at one time and survival assessed at a later time, necessarily reflecting survival consequences accumulated across the interval. The demographic processes operating within this interval are not directly observed; they constitute a demographic “black box” through which the ultimate conspecific density effects (and downstream coexistence outcomes) are realized (Fig. 1C) and statistically summarized (Fig. 1E). However, these demographic processes must be considered to fully understand what may drive variation in this accumulated CNDD. We argue that features of juvenile demography — namely density-independent mortality and growth rates — cause CNDD to vary in ways currently unrecognized.

This gap becomes evident when considering why CNDD varies across abiotic gradients. Empirically, CNDD is often stronger in wetter conditions (Swinfield et al., 2012; LaManna et al., 2016; Magee et al., 2021; Zang et al., 2021; Lebrija-Trejos et al., 2023; Comita et al., 2014; Bachelot et al., 2015; Uriarte et al., 2018; Song et al., 2018; Milici et al., 2020). CNDD variation has also been linked to elevation, soil chemistry, light availability, nutrient status, climatic variability, and habitat fragmentation (e.g., Fibich et al., 2021; LaManna et al., 2022; Xu et al., 2022; LaManna et al., 2016, 2017; Inman-Narahari et al., 2016; McCarthy-Neumann and Ibáñez, 2013; McCarthy-Neumann and Kobe, 2019; Song et al., 2021*b*; Holík et al., 2021; Magee et al., 2021; Zang et al., 2021; Huang et al., 2020; Krishnadas et al., 2018). These patterns are typically attributed to abiotic conditions directly modifying biotic interactions and thus altering instantaneous density sensitivity (i.e., affecting *α* in Fig. 1D). For example, LaManna et al. (2016) argue that resource-rich environments may strengthen CNDD by directly increasing host-specific natural enemy pressure or sensitivity to intraspecific resource competition. Similarly, recent work has suggested that higher precipitation and humidity may strengthen CNDD by enhancing phytopathogen transmission and host specificity of moisture-sensitive natural enemies (Milici et al., 2020; Lebrija-Trejos et al., 2023; Krishnadas, 2023; Milici, 2024). Alternatively, we hypothesize that abiotic gradients could alter realized interval CNDD by changing density-independent demography rather than instantaneous density sensitivity (Fig. 1D), and consequently be reflected in measured CNDD (Fig. 1E).

A similar issue arises when considering why CNDD varies among species. Empirically, species often differ substantially in measured CNDD (e.g., Klironomos, 2002; Petermann et al., 2008; Comita et al., 2010; Mangan et al., 2010; Johnson et al., 2017; Murphy et al., 2017; Song et al., 2021*a*; Hülsmann et al., 2024) (Fig. 1E). In particular, interspecific variation in measured CNDD often correlates with density-independent vital rates such as growth rate (faster-growing species exhibit stronger CNDD; Zhu et al., 2018; Song et al., 2021*b*; Qin et al., 2022; Huanca-Nunez et al., 2026). These patterns are usually interpreted as evidence that species differ in susceptibility to natural enemies or in sensitivity to intraspecific competition, and thus differ in instantaneous density sensitivity (*α* in Fig. 1D) (LaManna et al., 2024). However, we hypothesize that interspecific differences in density-independent vital rates could themselves generate variation in measured CNDD, independent of any difference in instantaneous density sensitivity.

We develop simple demographic models to elucidate how variation in juvenile demography may shape observed patterns of CNDD variation, and how that variation is reflected in empirical estimates. We ask three questions. First, can density-independent vital rates alter the survival consequences of conspecific crowding independently of instantaneous density sensitivity? Second, how does the demographic pathway through which density acts – for example, mortality versus growth, or effects concentrated in smaller versus larger individuals – shape the strength and direction of these effects? Third, could these demographic effects be reflected in interspecific and environmental variation in measured CNDD?

## Material and methods

We quantify CNDD using census-style survival summaries comparable to those used in empirical analyses, using a minimal pathogen model as a mechanistic proof of concept and a size-structured seedling model to examine more complex demographic scenarios. For each model, we simulate the corresponding stochastic process using a Gillespie Stochastic Simulation Algorithm (SSA) (Gillespie, 1977), generating individual-level census data. We analyze both a closed-cohort variant, in which no new individuals enter after the cohort is established, and an open, equilibrium variant, in which standing density is maintained by ongoing recruitment.

### Pathogen model

We consider a continuous-time stochastic Susceptible–Infected (SI) model of pathogen infection in a seedling cohort with host mortality. Susceptible individuals die at baseline rate *m*; infected individuals experience additional mortality *m*_inf_ (pathogen virulence). Susceptible individuals become infected via baseline exposure (rate *ξS*, representing spontaneous infection) or via mass-action transmission (rate *βSI*). The population receives a constant seedling supply at rate *f* . The mean-field equations are

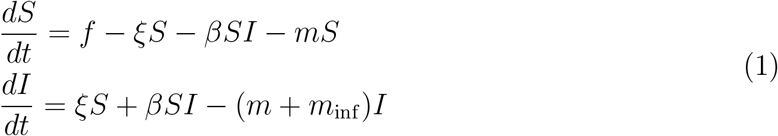

The closed-cohort variant (setting *f* = 0 and initiating the model with a cohort of susceptible seedlings *S*(0)) is appropriate for seedlings in their first year of life after a seasonal reproductive pulse (e.g., Lebrija-Trejos et al., 2023); the equilibrium variant (*f >* 0) is appropriate for established populations at later seedling stages with ongoing recruitment (e.g., Comita et al., 2023). For each variant, we analyzed how baseline mortality rate *m* influences CNDD.

For the closed-cohort variant, we used a Gillespie SSA corresponding to Eq. 1, where events occur at rates *λ*_infection_ = *ξS* + *βSI, λ*_death,*S*_ = *mS*, and *λ*_death,*I*_ = (*m* + *m*_inf_ )*I*. Each replicate began with an entirely susceptible cohort, *S*(0) = *N* (0) and *I*(0) = 0, and was run until a fixed census time *T*, after which we recorded *N* (*T* ) = *S*(*T* ) + *I*(*T* ) and the proportion that survived *N* (*T* )*/N* (0). We ran simulations varying *N* (0) and *m* while holding (*ξ, β, m*_inf_ ) fixed. For each (*N* (0), *m*) combination, we ran 20000 independent replicate simulations. To quantify how *m* shapes measured CNDD, we fit a separate Generalized Linear Model (GLM) for each value of *m*, modeling the survival outcome *y*_*k*_ ∈ *{*0, 1*}* of replicate *k* as

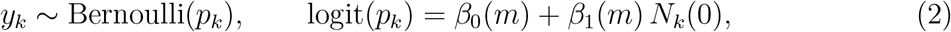

where more negative *β*_1_(*m*) indicates stronger CNDD. Note, unfortunately, *β* in the SI model and *β*_*i*_(*m*), *i* = 0, 1 share overlapping notation – we maintain this because they are standard in their fields.

We performed a similar analysis for the equilibrium variant. For each value of *m*, we selected *f* values to target a range of equilibrium densities *N*^∗^, allowing us to isolate the effect of *m* on CNDD independently of any change in equilibrium density. For each (*m, N*^∗^) combination, we used a Gillespie SSA with burn-in to reach the steady state, then tracked the fate of the cohort present at the end of the burn-in over a census interval *T*, and fit Eq. 2 to recover 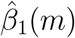 as a function of *m*.

### Size-structured seedling model

We use a coupled seed–seedling partial differential equation (PDE) model. Seeds are tracked as a local seed pool *B*(*t*), and seedlings are tracked as a size-structured cohort with density *n*(*x, t*) on *x* ∈ [0, *x*_max_], where *n*(*x, t*) *dx* is the number of individuals with size in [*x, x* + *dx*] at time *t*.

Seed dynamics follow

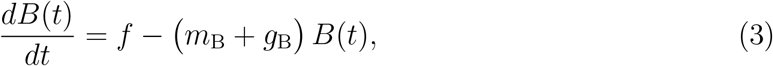

where *f* is seed input rate, *m*_B_ is seed loss rate, and *g*_B_ is germination rate. The closed cohort variant assumes *f* = *g*_B_ = 0 (ignoring seeds); the equilibrium variant assumes *f, g*_B_ *>* 0.

Seedlings follow an advection–mortality PDE analogous to the McKendrick–von Foerster equation (McKendrick, 1926; von Foerster, 1959) applied to size

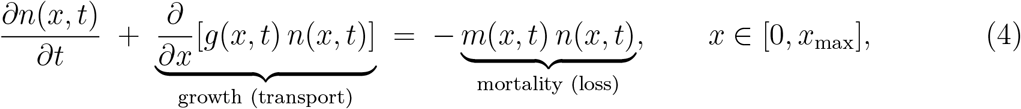

with recruitment from the seed pool entering as an inflow boundary condition at the smallest size *g*(0, *t*) *n*(0, *t*) = *g*_B_ *B*(*t*). The closed cohort model assumes all seedlings begin at size *x* = 0.

Mortality rate is specified as a function that decreases with size *x* (as observed empirically) and increases with density (*α >* 0):

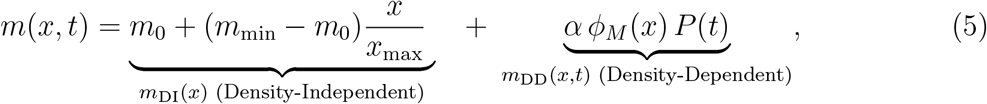

where *m*_0_ is the hazard at the smallest size, *m*_min_ is the hazard at *x* = *x*_max_; see below for density-dependent terms. Growth rate is given by

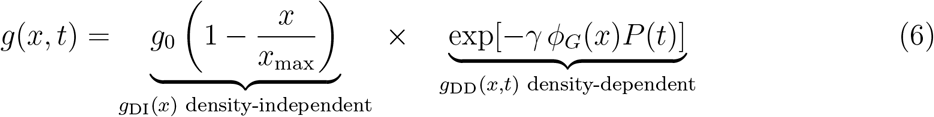

where *g*_0_ is baseline growth rate; individuals grow toward maximum size *x*_max_.

The density-dependent terms in Eqs. 5–6 depend on *α* and *γ*, which represent the *instantaneous density sensitivity* of mortality and growth, respectively (Fig. 1D). Crowding effects depend on both the size distribution of neighbors and the size-dependent susceptibility of the focal size class. The neighborhood crowding index

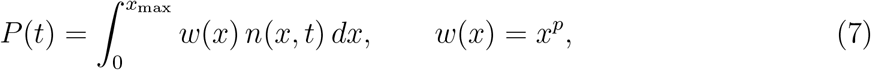

captures how the size of neighboring individuals weights their contribution to crowding: *p* = 0 corresponds to abundance weighting (size-independent effects), *p >* 0 upweights larger neighbors, and *p <* 0 upweights smaller neighbors. Size-dependent susceptibility of the focal size class is captured by *ϕ*_*M*_ (*x*) and *ϕ*_*G*_(*x*), each of the form

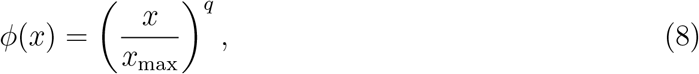

where *q* = 0 implies no size dependence in susceptibility to density effects, *q <* 0 implies stronger effects on smaller individuals, and *q >* 0 implies stronger effects on larger individuals.

#### Spatially explicit multispecies simulations

We simulated a multispecies, equilibrium-variant version of the size-structured model using a Gillespie SSA, generating census-style survival datasets. We used these simulations for two purposes: first, to test how spatial environmental heterogeneity affecting density-independent vital rates can induce environment-by-density interactions in fitted survival models; second, to study how interspecific differences in density-independent vital rates map onto interspecific variation in CNDD.

Simulations were run on a grid of discrete patches (Fig. 1A). Each patch was assigned an environmental value *E*_*j*_ drawn at initialization with specified spatial autocorrelation (via spatial smoothing) and then held fixed through time. Adults were placed on the grid and held static throughout each simulation — no adult mortality or recruitment — with uneven abundances across species (lognormal draws) and spatial aggregation generated by species-specific autocorrelated surfaces.

Each adult produces seeds at rate *f* and disperses them according to a Gaussian dispersal kernel. Seed loss events remove one seed at constant per-capita rate *m*_*B*_, and germination events (rate *g*_*B*_) convert a seed into a seedling placed in the smallest size bin. Seedlings are tracked as integer abundances across discrete size bins. Size is discretized on [0, *x*_max_] into *b* equal bins of width Δ*x* = *x*_max_*/b*, with bin *ℓ* spanning [(*ℓ* − 1)Δ*x, ℓ*Δ*x*] and midpoint 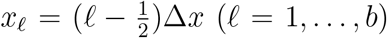 individuals in bin *ℓ* are treated as located at *x*_*ℓ*_. We track abundances *N*_*s,ℓ*_(*t*), where *s* indexes species and *ℓ* indexes size bins. Growth is implemented as transitions from bin *ℓ* to bin *ℓ*+1 at per-capita rate *g*(*x*_*ℓ*_, *t*)*/*Δ*x*, and mortality as per-capita deaths from bin *ℓ* at rate *m*(*x*_*ℓ*_, *t*). The mean-field limit corresponds to the finite-volume approximation of Eq. 4 (Supplementary Information Section 9).

The multispecies Gillespie SSA simulations distinguish two sources of crowding for each focal species *s*: conspecific crowding and heterospecific crowding. Both follow the same size-weighting scheme as before (*w*(*x*) = *x*^*p*^, Eq. 7). For an individual *i* of species *s*(*i*) in size bin *ℓ*(*i*), conspecific crowding is *P*_con,*s*(*i*)_(*t*) = ∑_*ℓ*_ *w*(*x*_*ℓ*_) *N*_*s*(*i*),*ℓ*_(*t*) − *w*(*x*_*ℓ*(*i*)_), excluding the individual’s own contribution to bin *ℓ*(*i*); heterospecific crowding is pooled across all other species, *P*_het,*s*(*i*)_(*t*) = ∑_*k*≠*s* (*i*)_ ∑_*ℓ*_ *w*(*x*_*ℓ*_) *N*_*k,ℓ*_(*t*). These crowding components enter the mortality and growth functions (Eqs. 5–6) additively, replacing the *P* (*t*) term, with species-specific sensitivity parameters *α*_con,*s*_, *α*_het,*s*_ (mortality) and *γ*_con,*s*_, *γ*_het,*s*_ (growth) governing sensitivity to each crowding source. *ϕ*_*M*_ (*x*_*ℓ*_) and *ϕ*_*G*_(*x*_*ℓ*_) (Eq. 8) govern how susceptibility varies with size.

For our first purpose (environmental effects on CNDD), we allowed a vital rate component *θ* (*m*_0_ or *g*_0_) to vary across patches. Patch environments *E*_*j*_ were constructed as a Gaussian-smoothed i.i.d. *N* (0, 1) field, standardized to mean 0 and unit variance. We implemented environmental effects as *θ*(*E*_*j*_) = *θ*_baseline_ exp(*η*_*θ*_*E*_*j*_), in which the baseline value of the vital rate *θ*_baseline_ was modified by the factor exp(*η*_*θ*_*E*_*j*_) through space. We then fit survival models (see below) to examine the resulting environment-CNDD interactions.

For our second purpose (interspecific CNDD variation), we drew vital-rate parameters (*m*_0_ and *g*_0_) independently from lognormal distributions for each species and compared fitted species-level CNDD estimates (see next section) to the vital-rate values used to generate each species’ dynamics.

Full parameter values used in simulations are given in Tables S2–S3. Supplementary Information section 10 provides additional details on simulations and model fits described below.

#### Statistical models for simulated censuses

From Gillespie simulations, we constructed census windows of length *T* = 5 time units over 350 total time units on 800 patches. This yielded between 5 *×* 10^5^ and 10^6^ individual-level survival records, forming the dataset for Generalized Linear Mixed Model (GLMM) fitting. Since the Gillespie SSA evolves binned counts *N*_*s,ℓ*_(*t*) rather than individual trajectories, we reconstructed individual fates by assigning each seedling a unique ID at germination and, when a growth or death event occurred in bin (*s, ℓ*), selecting one of the *N*_*s,ℓ*_(*t*) individuals at random to enter the next size bin or die.

For each census window, let *y*_*i*_ = 1 if seedling *i* is alive at the end of the interval and *y*_*i*_ = 0 otherwise. Following typical methods of CNDD estimation, we modeled survival with a Bernoulli GLMM with a logit link. For seedling *i* of species *s*(*i*), let *p*_*i*_ = Pr(*y*_*i*_ = 1) denote survival probability; we modeled

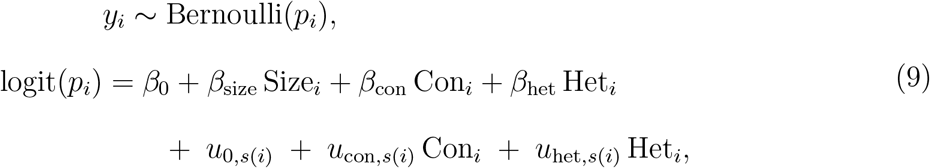

where Con_*i*_ and Het_*i*_ are the conspecific and heterospecific crowding covariates at the initial census, and Size_*i*_ is initial size. The terms (*u*_0,*s*_, *u*_con,*s*_, *u*_het,*s*_) are species-level random effects. *β*_con_ and *u*_con,*s*_ are our estimates of measured CNDD at the population and species levels, respectively.

In simulations with spatial environmental heterogeneity, we modified Eq. 9 by adding fixed effects of patch environment and environment-by-Con-density interactions *β*_*E*_ *E*_*j*(*i*)_ + *β*_*E*:con_ *E*_*j*(*i*)_ Con_*i*_ where *E*_*j*(*i*)_ is the environment of the patch containing individual *i. β*_*E*:con_ captures how strongly conspecific density effects on survival vary with the environment.

We accounted for nonlinearities in neighborhood effects (Detto et al., 2019) by applying a power transform to raw crowding measures: 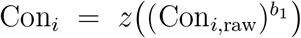 and 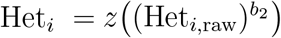, where Con_*i*,raw_ and Het_*i*,raw_ are conspecific and heterospecific neighbor densities, respectively, and *z*(·) denotes z-scoring within the fitted dataset (with Size, and when included *E*, standardized analogously). The exponents (*b*_1_, *b*_2_) were selected by a grid search over candidate values with *b*_1_, *b*_2_ ∈ *{*0.25, 0.5, 0.75, 1, 1.25, 1.5, 2*}*, choosing the combination that maximized the fitted log-likelihood for each fit. For robustness, we performed analogous analyses using Generalized Additive Models (GAMs).

Simulations were performed in Julia (v1.11.6) (Bezanson et al., 2017). Statistical analyses were performed in `R` (v4.3.2) (R Core Team, 2022) using `lme4` (Bates et al., 2015) for GLMMs and `mgcv` (Wood and Wood, 2015) for GAMs.

### Hazard-based metrics and connection to CNDD estimates

Seedling survival is the central quantity of interest in both the pathogen model (Eq. 1) and the size-structured model (Eq. 4). We connect both models to empirical CNDD estimates using the mortality hazard function *h*(*t*). For an individual alive at time *t*, the probability of dying in [*t, t* + *dt*] is *h*(*t*) *dt* on average; when individuals differ in state (e.g., size, infection), *h*(*t*) denotes the cohort-averaged hazard. For a cohort with no recruitment during [0, *T* ], abundance declines as *dN/dt* = −*h*(*t*) *N* (*t*), giving

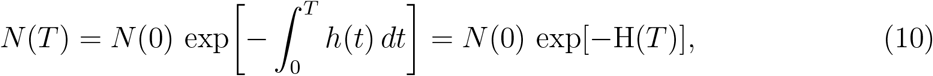

where 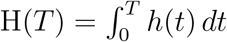 is the cumulative hazard over [0, *T* ], and the cohort survival fraction is *N* (*T* )*/N* (0) = exp[−H(*T* )].

For each model, we write the cohort-averaged hazard as 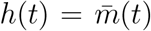. In the pathogen model, 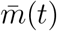 is the baseline mortality rate plus the additional infection-driven mortality, weighted by the proportion of the cohort currently infected:

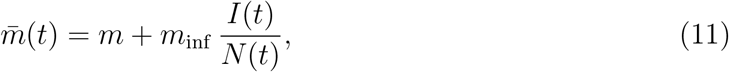

where *N* (*t*) = *S*(*t*) + *I*(*t*). In the size-structured PDE model, 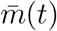 is instead an average over sizes,

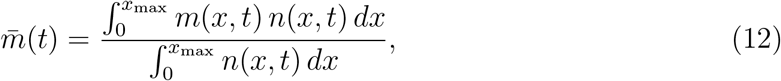

where *m*(*x, t*) is given by Eq. 5 and 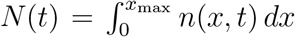 is the total density. For both models, the excess mortality attributable to density dependence (Fig. 1D) is

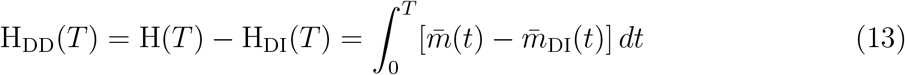

where 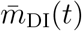 is the cohort-averaged density-independent hazard from a matched simulation with density dependence removed (transmissibility *β* = 0 for the pathogen model; *α* = *γ* = 0 for the size structured model), and 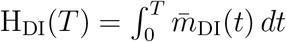 is its cumulative counterpart. We interpret H_DD_(*T* ) as *realized interval CNDD*.

The key quantity CNDD studies aim to capture is how survival probability exp (−H(*T* )) responds to initial conspecific density *N* (0). The statistical models described above (Eqs. 2, 9) are common instances of this approach: modeling the survival probability of individual *i, p*_*i*_, as logit(*p*_*i*_) = *β*_0_ +*β*_1_ *N* (0)+…, where *N* (0) is the initial conspecific density *i* experiences. More negative *β*_1_ indicates stronger CNDD. We assume this logit-linear form for tractability. Since *p*_*i*_ = exp(−H(*T* )), differentiating logit(*p*_*i*_) with respect to *N* (0) gives the local logit-scale density effect:

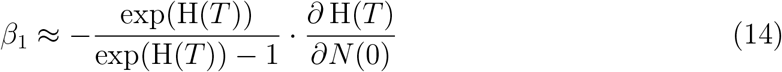

(see Supplementary Information Section 1). Eq. 14 is exact when logit-survival is linear in *N* (0) and otherwise gives the local logit-scale density effect that an estimate of *β*_1_ approximates as a weighted average over the observed density range.

Since H_DI_(*T* ) is independent of density *N* (0), *∂*H(*T* )*/∂N* (0) = *∂*H_DD_(*T* )*/∂N* (0): the sensitivity of cumulative hazard to initial density equals the sensitivity of realized interval CNDD to initial density. Measured CNDD, *β*_1_, is thus proportional to the density-sensitivity of realized interval CNDD; we use this property in our analyses.

### Presentation of results

First, we examine how baseline mortality shapes pathogen-mediated CNDD, analyzing Eq. 1. Second, we derive general relationships between density-independent vital rates and CNDD, analyzing the size-structured model (Eq. 4). Third, we use the multispecies Gillespie SSA and GLMMs (Eq. 9) to examine how spatial variation in baseline vital rates generates environment-dependent measured CNDD, and how interspecific variation in vital rates influences interspecific variation in measured CNDD. Throughout, we use the hazard-based approach to evaluate how vital rates influence CNDD strength, *β*_1_.

## Results

### CNDD decreases with increasing density-independent mortality in the pathogen model

Baseline mortality *m* weakens pathogen-mediated CNDD by suppressing infection outbreaks in closed seedling cohorts (*f* = 0). At low initial density, infection rarely establishes regardless of *m*, so the excess per-capita mortality rate due to infection, *m*_inf_ *I*(*t*)*/N* (*t*), stays negligible (Fig. 2A,C). At high initial density, lower baseline mortality allows infections to establish early and spread extensively, producing a substantially larger accumulated mortality from infection — realized interval CNDD (H_DD_(*T* ), Eq. 13) — over interval [0, *T* ] (Fig. 2C, “low base mortality,” orange trail segments; Fig. 2D, blue/purple curves); higher *m* suppresses this accumulation, as hosts die before infections can establish and transmit (Fig. 2C, “high base mortality”; Fig. 2D, yellow/green curves). Correspondingly, the probability of an infection outbreak decreases with *m* (Fig. 2E), and measured CNDD weakens accordingly: 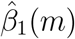 becomes progressively less negative and approaches zero at high *m* (Fig. 2F,G). An approximation of *β*_1_(*m*), derived from Eq. 14, highlights its dependence on *m*:

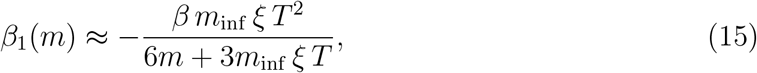

which closely tracks the GLM-fitted 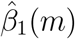 (Fig. 2I; Supplementary Information Section 3). Biological intuition is provided by approximating early pathogen spread as a birth–death process (see Supplementary Information Section 2 for derivations). A single infected seedling produces new infections at rate *βS*(*t*) and dies at rate *m* + *m*_inf_ ; their ratio gives the effective reproduction number *R*_*t*_ = *βS*(*t*)*/*(*m* + *m*_inf_ ). Susceptibles decline prior to any outbreak such that *S*(*t*) ≈ *N* (0)*e*^−*mt*^, and *R*_*t*_ ≈ *R*_0_ *e*^−*mt*^, where *R*_0_ ≡ *βN* (0)*/*(*m* + *m*_inf_ ) is the basic reproduction number. Baseline mortality *m* enters *R*_*t*_ by (i) thinning the susceptible pool over time (*e*^−*mt*^) and (ii) reducing *R*_0_ directly. Infection can only establish while *R*_*t*_ *>* 1, and this time window shrinks with *m*. Introductions occur at rate *λ*_intro_(*t*) ≈ *ξS*(*t*), with each introduction succeeding with probability *q*(*t*) ≈ max*{*0, 1 − 1*/R*_*t*_*}*. Successful establishments form a nonhomogeneous Poisson process with intensity *r*(*t*) = *λ*_intro_(*t*) *q*(*t*), yielding pathogen establishment probability

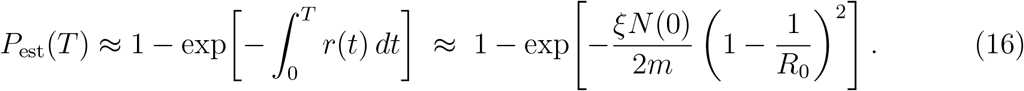

*P*_est_(*T* ) declines with *m* (Fig. 2E).

**Figure 2.**
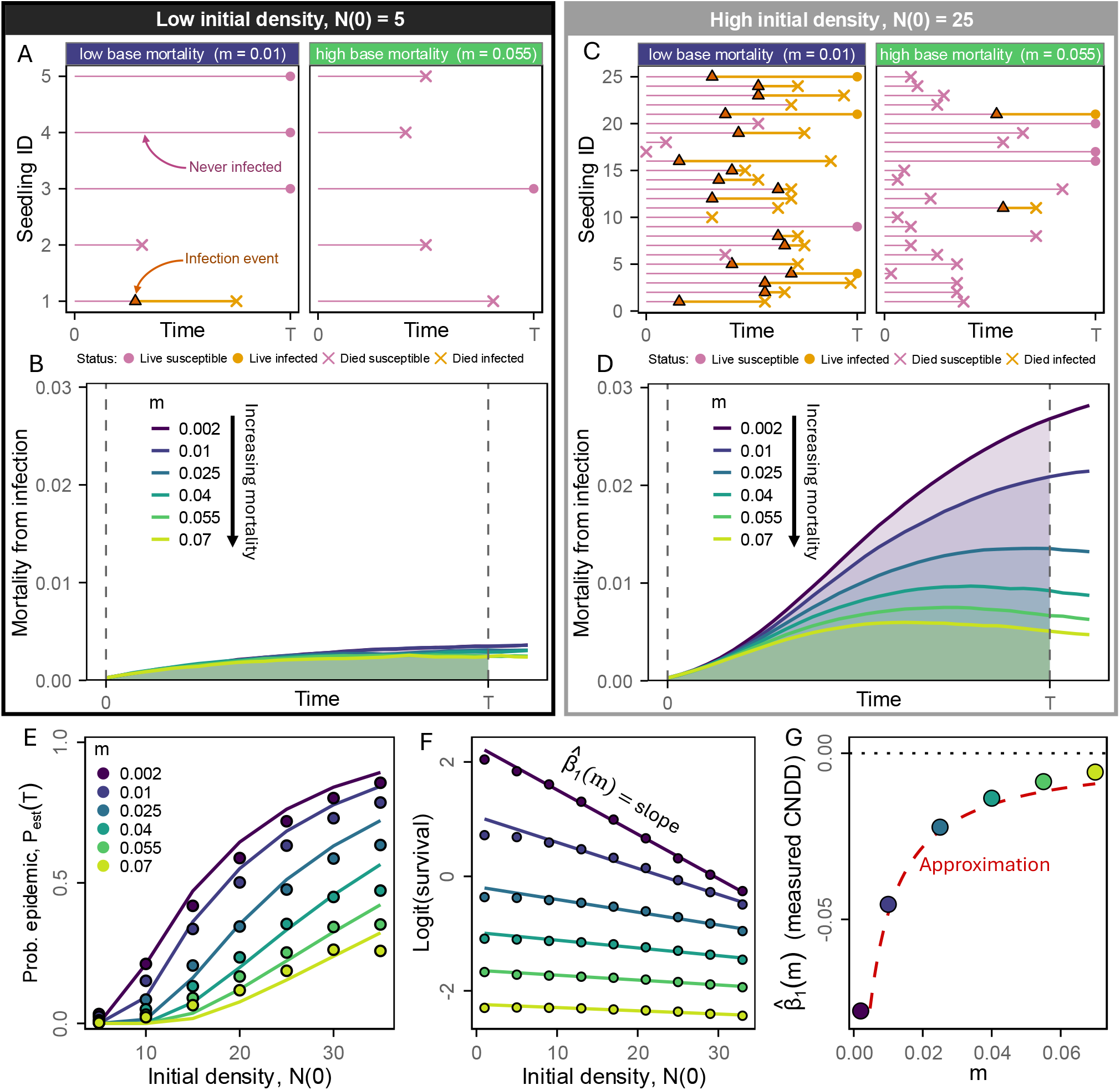
Density-independent mortality weakens pathogen-mediated CNDD by suppressing infection establishment. (A, C) Example Gillespie simulation realizations of the closed cohort stochastic susceptible–infected (SI) model showing individual infection and mortality event histories at low initial density (*N* (0) = 5; A) and high initial density (*N* (0) = 25; C), for low baseline mortality (*m* = 0.01; left panel) versus high baseline mortality (*m* = 0.055; right panel). Horizontal lines show individual lifespans: pink segments indicate the susceptible period; orange segments indicate the infected period following the onset of infection (filled triangle). Points indicate the final state of each individual at the census time *T* (filled circles: alive; X’s: dead; colors indicate infection status). (B, D) Mean mortality from infection *m*_DD_(*t*) = *m*_inf_ · *I*(*t*)*/N* (*t*) (Eq. 11) over the census interval [0, *T* ] (dashed lines), averaged over simulation replicates, for low initial density (B) and high initial density (D), shown for each value of baseline mortality *m*. Shaded area shows cumulative density-dependent mortality — realized interval CNDD H_DD_(*T* ) (Eq. 13) — over [0, *T* ]. (E) Probability of infection establishment (an epidemic) *P*_est_(*T* ) as a function of cohort size *N* (0) for different *m*. Curves show analytical approximations; points show simulation means (establishment is defined as ≥ 10% of the initial cohort becoming infected during [0, *T* ]). (F) Logit survival as a function of *N* (0) for each *m*. Points are calculated from simulation means; lines are fitted binomial GLMs for each *m*. (G) Measured CNDD 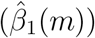 as a function of *m*. More negative values indicate stronger CNDD; red dashed line is the approximation (Eq. 15). Parameters: *ξ* = 0.0025, *β* = 0.015, *m*_inf_ = 0.1, *T* = 33.

Results are qualitatively identical for the equilibrium variant (*f >* 0; Supplementary Information Section 4; Fig. S1). Lower baseline mortality *m* sustains a higher fraction of infected individuals at any given equilibrium density *N*^∗^, so infection-driven mortality increases more steeply with conspecific density when *m* is lower (Fig. S1A-E). GLM fits to census data generated from equilibrium simulations yield 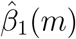 values that decrease with *m* (Fig. S1F).

### How do density-independent vital rates shape CNDD?

We now analyze the size-structured model (Eq. 4). We first consider how density-independent vital rates shape the density-dependence of survival within closed cohorts (*f* = *g*_B_ = 0) of seedlings starting in the smallest size class. Unsurprisingly, greater instantaneous sensitivity to density (*α* or *γ*) steepens this density-dependence (Box 1; Fig. S2). Less trivially, baseline growth (*g*_0_) and mortality (*m*_0_) can amplify or attenuate this density-dependence when instantaneous density sensitivity (*α* or *γ*) is held fixed. Whether changes in these parameters strengthen or weaken the density-dependence of survival depends on (i) which vital rate density affects (mortality vs. growth), and (ii) which size classes generate and/or experience the density-dependent feedbacks. Below, we outline four (nonexhaustive) characteristic cases (Fig. 3; Box 1).

**Figure 3.**
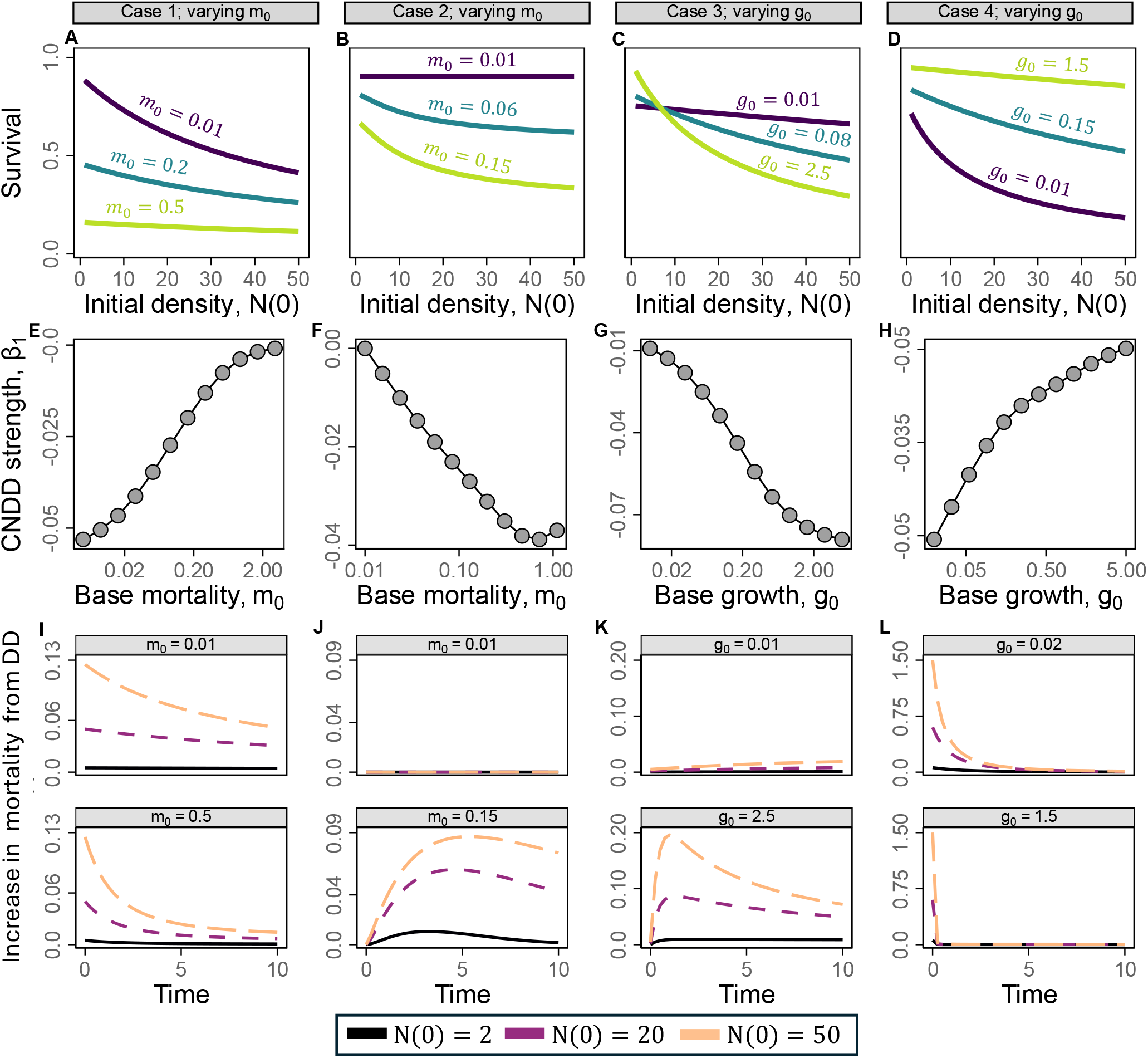
Density-independent vital rates modulate CNDD. Each column shows a distinct demographic scenario (Cases 1–4, as described in Box 1 and the main text). All panels are generated by numerically solving the full size-structured PDE model (Eq. 4, with vital rates given by Eqs. 5–6). (A–D) Survival probability exp( −H(*T* )) = *N* (*T* )*/N* (0) at interval time *T* as a function of initial cohort size *N* (0), for the labeled ranges of focal density-independent parameters in each panel. (E–H) Measured CNDD strength, summarized as *β*_1_ — the exact logit-scale density slope (Eq. 14) — as a function of the focal parameter; more negative values indicate stronger CNDD. *β*_1_ is computed by evaluating *∂*H(*T* )*/∂N* (0) via central difference at each *N* (0) on a grid from 1 to 50 and averaging across that grid. (I–L) The excess instantaneous per-capita hazard attributable to density dependence 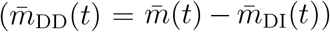 over the census interval, shown for a low (top facet) and high (bottom facet) value of the focal parameter, and for three initial cohort sizes (line colors; see legend); the area under each curve is realized interval CNDD, H_DD_(*T* ) (Eq. 13). Columns correspond to: (A,E,I) varying baseline small-size mortality *m*_0_ when density acts through mortality (*α >* 0, *γ* = 0); (B,F,J) varying *m*_0_ when density acts through growth suppression (*γ >* 0, *α* = 0); (C,G,K) varying baseline growth rate *g*_0_ when density-dependent effects increase with individual size; and (D,H,L) varying *g*_0_ when density-dependent effects are concentrated in small individuals. All other parameters are held fixed within each column; see Supplementary Information Table S1 for full parameter list.

#### Case 1: density acts on mortality, symmetrically across sizes

Higher baseline mortality weakens the density-dependence of survival (Box 1, Case 1) when density dependence acts directly on mortality (*α >* 0, *γ* = 0), affecting all individuals regardless of size. Higher baseline mortality *m*_0_ lowers overall survival but flattens its dependence on initial density (Fig. 3A), and *β*_1_ (Eq. 14) becomes progressively less negative (weaker CNDD) as *m*_0_ increases (Fig. 3E). Conceptually, higher density-independent mortality means that individuals, on average, are both exposed to conspecific neighbors for less time — because neighbors die faster — and themselves accumulate less density-dependent hazard before dying. This reduces the opportunity for density-dependent effects to accumulate over [0, *T* ] (Fig. 3I). This resembles the SI model result.

#### Case 2: density suppresses growth, with mortality varying by size

In contrast to Case 1, when density dependence acts by reducing growth (*γ >* 0, *α* = 0), the strength of density-dependence depends on the mortality contrast between smaller and larger individuals (Box 1, Case 2). If small individuals experience higher baseline mortality than large individuals (*m*_0_ *> m*_min_) — typical for seedlings — then growth suppression retains individuals longer in the high-mortality stage. This retention increases cumulative hazard and steepens the density-dependence of survival. Hence, survival declines more steeply with higher *m*_0_ (Fig. 3B), and *β*_1_ becomes more negative (stronger CNDD) with increasing *m*_0_ (Fig. 3F). When mortality does not vary with size (*m*_0_ = *m*_min_), density-dependent growth suppression does not affect survival (Fig. 3J, top facet). Greater baseline growth rate *g*_0_ also strengthens CNDD in Case 2: growth suppression weakly affects slow growers (Supplementary Information Section 7, Fig. S3).

#### Case 3: large individuals generate most density-dependent pressure

Faster baseline growth *g*_0_ steepens the density-dependence of survival (Box 1, Case 3) when density-dependent pressure is generated predominantly by larger individuals (effects scale with size *x*; *p* = 1 in Eq. 7). Higher growth increases overall survival but steepens its dependence on initial density (Fig. 3C), and *β*_1_ becomes more negative as *g*_0_ increases (Fig. 3G). Biologically, faster growth produces large individuals more rapidly, increasing how much density-dependent mortality the cohort experiences over [0, *T* ] (Fig. 3K).

#### Case 4: small individuals both generate and experience density-dependent mortality

In contrast to Case 3, if density dependence is generated by and acts most strongly on small individuals (*q, p <* 0 in Eqs. 7 and 8), then faster baseline growth *g*_0_ flattens the density-dependence of survival (Fig. 3D). Accordingly, *β*_1_ becomes less negative as *g*_0_ increases (Box 1, Case 4; Fig. 3H). Biologically, baseline growth functions as an escape rate from the stage where density dependence is strongest: faster growth moves both focal individuals out of the vulnerable stage and neighbors out of the density-generating stage more rapidly, decreasing how much density-dependent mortality the cohort experiences over [0, *T* ] (Fig. 3L).

#### Equilibrium analysis

Analyses assuming *f >* 0 yield qualitatively identical results to the closed-cohort model in all four cases, though the underlying mechanisms differ slightly in detail. For example, in Case 3, increased CNDD from faster baseline growth *g*_0_ is driven not by a change in the sizes of the closed-cohort individuals over the census interval, but by higher *g*_0_ shifting the equilibrium size distribution *n*^∗^(*x*) toward larger individuals, steepening the density-dependence of survival since crowding is generated by larger individuals. See Supplementary Information Section 8 and Figs. S4–S5 for details.

##### Box 1. Two-stage approximation for *β*1 analysis

A central result of our analysis is that density-independent vital rates — baseline mortality and growth — shape the regression coefficient *β*_1_ estimated in empirical CNDD studies (Eq. 14), independently of instantaneous density sensitivity (*α* or *γ*). Here, we show this analytically. Under high survival, i.e. small cumulative hazard H(*T* ), Eq. 14 simplifies to

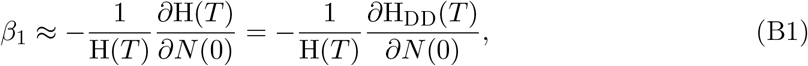

which we use throughout the four cases below (see Supplementary Information Section 1). We derive *β*_1_ analytically using a simplified two-stage approximation of the size-structured model (Eq. 4), collapsing it into a “small” stage with abundance *J*_1_(*t*) and a “large” stage with abundance *J*_2_(*t*), with individuals transitioning from small to large at per-capita rate *g*(*J*_1_, *J*_2_) and dying with stage-specific per-capita hazards *µ*_1_(*J*_1_, *J*_2_) and *µ*_2_(*J*_1_, *J*_2_):

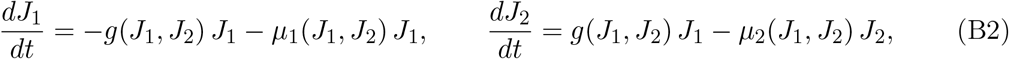

with *J*_1_(0) = *N* (0) and *J*_2_(0) = 0. Total cohort abundance is *N* (*t*) = *J*_1_(*t*) + *J*_2_(*t*). The cohort mean hazard — the average per-capita mortality rate — is 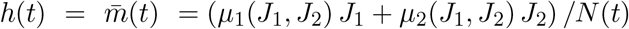, and the cohort cumulative hazard is 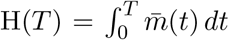. We derive Taylor expansions of H(*T* ) and *∂*H(*T* )*/∂N* (0) in *T*, retaining terms through second order (valid when effects are weak), and derive *β*_1_ via the above approximation. The four cases below illustrate how the demographic pathway through which density acts — whether through mortality or growth, and which size classes are involved — determines the specific way density-independent vital rates enter *β*_1_.

Results are qualitatively identical when instead examining the sensitivity of survival itself, *∂* exp( −H(*T* ))*/∂N* (0) ≈ −*∂*H(*T* )*/∂N* (0) = −*∂*H_DD_(*T* )*/∂N* (0) under high survival — the last equality following from H_DI_(*T* )’s independence from *N* (0) (Methods) — simply because both this quantity and *β*_1_ (Eq. B1) are proportional to *∂*H(*T* )*/∂N* (0). See Supplementary Information Section 5 for details and derivations.

**Case 1: Symmetric density-dependent mortality; density-independent transition**.

Assume density acts only through mortality, equally across stages, with constant transition rate:

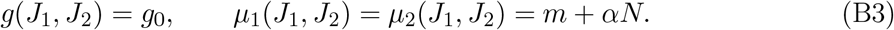

The corresponding regression coefficient is

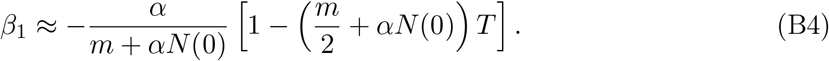

Higher baseline mortality *m* weakens CNDD (less negative *β*_1_) by shrinking the fraction of total mortality risk attributable to crowding, and by depleting conspecific density over the interval (Fig. 3A,E). Transition rate *g*_0_ has no effect, because stages are demographically identical.

**Case 2: Density suppresses growth/transition**. We assume mortality is density independent, but transition is crowding-suppressed:

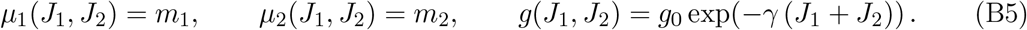

The corresponding regression coefficient is

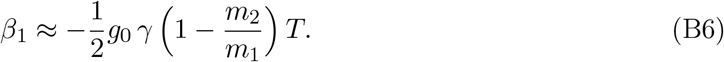

Growth suppression generates CNDD only if remaining in the small stage carries a mortality cost (*m*_1_ *> m*_2_). CNDD strengthens with a larger mortality contrast between stages (Fig. 3B,F) and with faster baseline growth *g*_0_ (Supplementary Information Fig. S3). Because density acts through growth suppression rather than mortality directly, CNDD builds up gradually over the interval rather than acting immediately, growing in magnitude with *T* (hence, *β*_1_ → 0 as *T* → 0).

**Case 3: Large individuals drive density-dependent mortality of small individuals**. Assume constant transition and density acts by increasing small-stage mortality in proportion to *J*_2_:

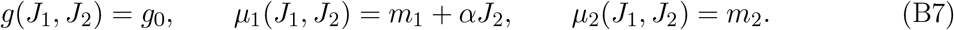

The corresponding regression coefficient is

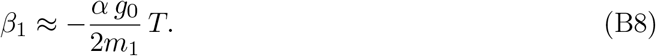

Like Case 2, this effect also builds up gradually: because *J*_2_(0) = 0, density-dependent mortality cannot act until large individuals accumulate. Consequently, *β*_1_ ∝ *T* . Faster transition *g*_0_ strengthens CNDD by accelerating this accumulation, while higher baseline small-stage mortality *m*_1_ weakens *β*_1_, since small individuals die before large individuals have time to accumulate (Fig. 3C,G).

**Case 4: Only small individuals generate and experience density-dependent mortality**. Assume constant transition and density dependence acts only on the small-stage hazard via *J*_1_:

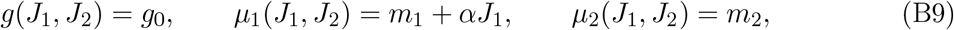

which is biologically feasible if, for example, pathogen transmission occurs primarily among small seedlings. The corresponding regression coefficient is

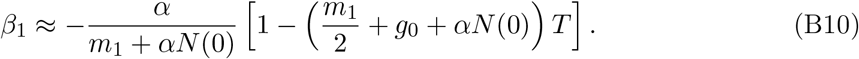

As in Case 1, higher *m*_1_ weakens *β*_1_. Unlike Cases 2 and 3, faster growth *g*_0_ weakens *β*_1_: larger *g*_0_ causes individuals to more rapidly exit the small size class, which both generates and suffers density-dependent effects (Fig. 3D,H).

### Why might CNDD vary on environmental gradients?

The demographic mechanisms highlighted in Cases 1–4 also generate environment-dependent measured CNDD. We tested this using our spatially-explicit simulations, varying density-independent vital rates across patches and fitting an environment-augmented GLMM (Eq. 9 with environment terms added; Fig. 4). Spatial variation in these vital rates alone — with *α* or *γ* held constant through space — was sufficient to produce significant Env*×*Con interactions (*β*_*E*:con_ ≠ 0; Fig. 4A,C,E,G), with the sign depending on the demographic scenario.

**Figure 4.**
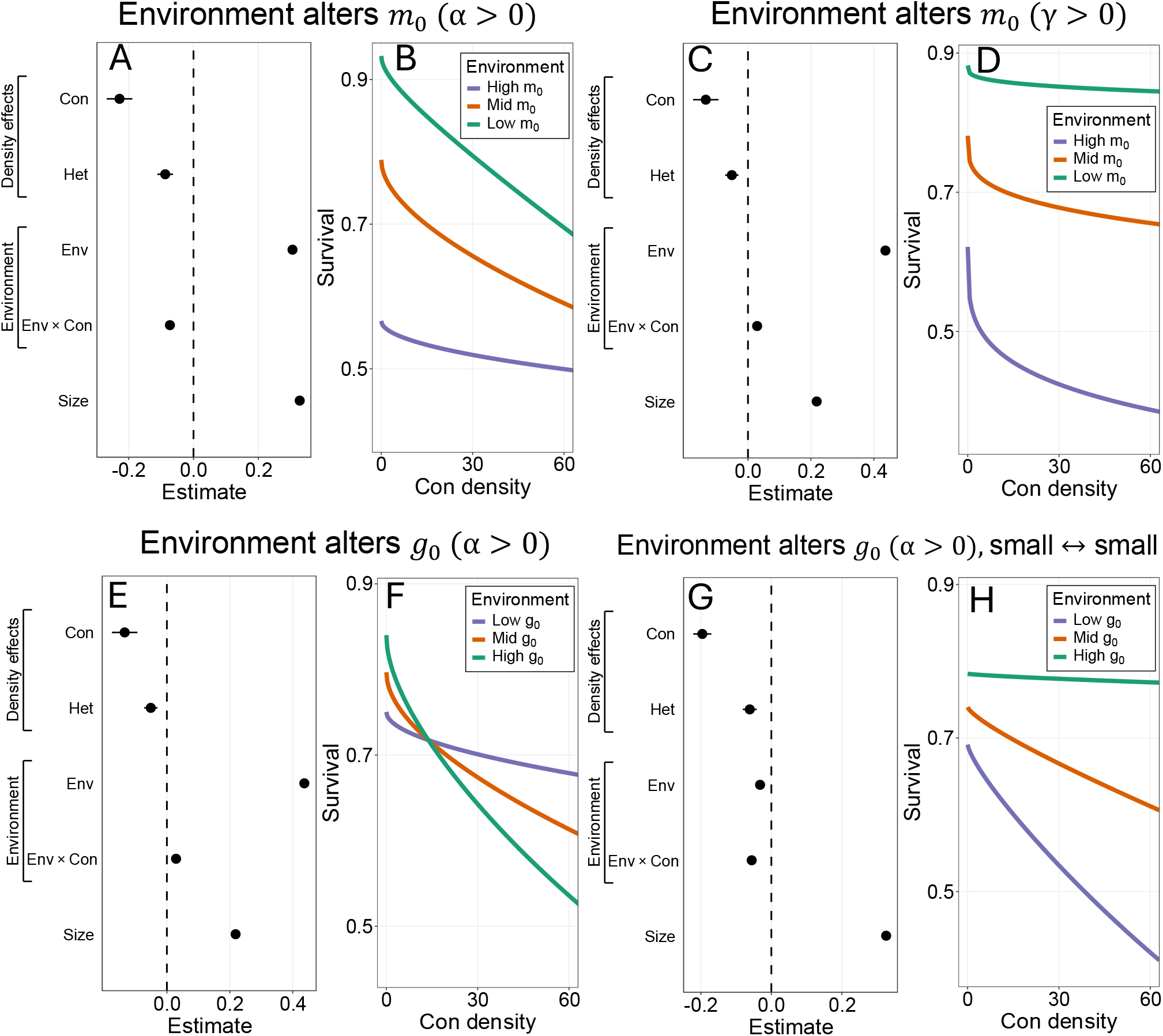
Density-independent vital rates mediate environment-dependence in measured CNDD. Panels summarize generalized linear mixed models (GLMMs) fit to census survival outcomes from spatially explicit simulations in which the environment varies among patches and affects a density-independent vital rate. GLMMs use the environment-augmented version of Eq. 9. (A,C,E,G) Standardized fixed-effect estimates (points with uncertainty intervals) for conspecific density (Con), heterospecific density (Het), the environmental covariate (Env), interactions between the environment and conspecific effects Env × Con, and size. (B,D,F,H) Population-level survival predictions from the fitted GLMMs as a function of conspecific density, evaluated at low, median, and high environments (1st, 50th, and 99th percentiles), with other covariates held at their means. (A,B) Akin to Case 1: environment alters small-seedling mortality *m*_0_ when density acts through mortality (*α >* 0, *γ* = 0). (C,D) Akin to Case 2: environment alters small-seedling mortality *m*_0_ when density acts through growth suppression (*γ >* 0, *α* = 0). (E,F) Akin to Case 3: environment alters baseline growth rate *g*_0_ when density acts through mortality (*α >* 0, *γ* = 0), with density feedback dominated by larger individuals. (G,H) Same as (E,F), but – akin to Case 4 – density effects are concentrated in small individuals. Simulations contain 75 species and included data generated from 800 patches on a 10,000 patch landscape. Full simulation and fitting details are provided in the Supplementary Information section 10; see Table S2 for all simulation parameters.

When the environment alters baseline mortality (*m*_0_) and density dependence acts through mortality (*α >* 0), lower-*m*_0_ environments show higher survival at low density but a steeper decline as density increases (Fig. 4A–B; Case 1). When density dependence instead acts through growth suppression (*γ >* 0), lower-*m*_0_ environments again show higher low-density survival, but now with a flatter decline — the reverse of Case 1 (Fig. 4C–D; Case 2). When the environment instead alters baseline growth rate (*g*_0_), faster-growth environments produce a steeper decline when larger individuals generate crowding (Fig. 4E–F; Case 3), but a flatter decline when crowding effects are concentrated among small individuals (Fig. 4G–H; Case 4).

### What regulates interspecific variation in measured CNDD?

Fitting GLMMs to data from our spatially-explicit simulations in which density-independent vital rates vary across species yields substantial interspecific variation in CNDD, quantified by species-specific random-slope estimates for conspecific density (Fig. 5). This variation follows the mechanisms established in Box 1 and Fig. 3 (Cases 1–4): species whose vital rates (*m*_0_, *g*_0_) steepen the density-dependence of survival exhibit stronger measured CNDD, and vice versa (Fig. 5A–H). Simulating conditions in which *m*_0_, *g*_0_, both, or neither varied across species (with identical *α >* 0; *γ* = 0), interspecific variation in measured CNDD was near zero only when neither vital rate varied, and substantial whenever either or both did (Fig. 5I). Results are qualitatively identical fitting GAMs to the simulated data (Supplementary Information section 11; Fig. S6).

**Figure 5.**
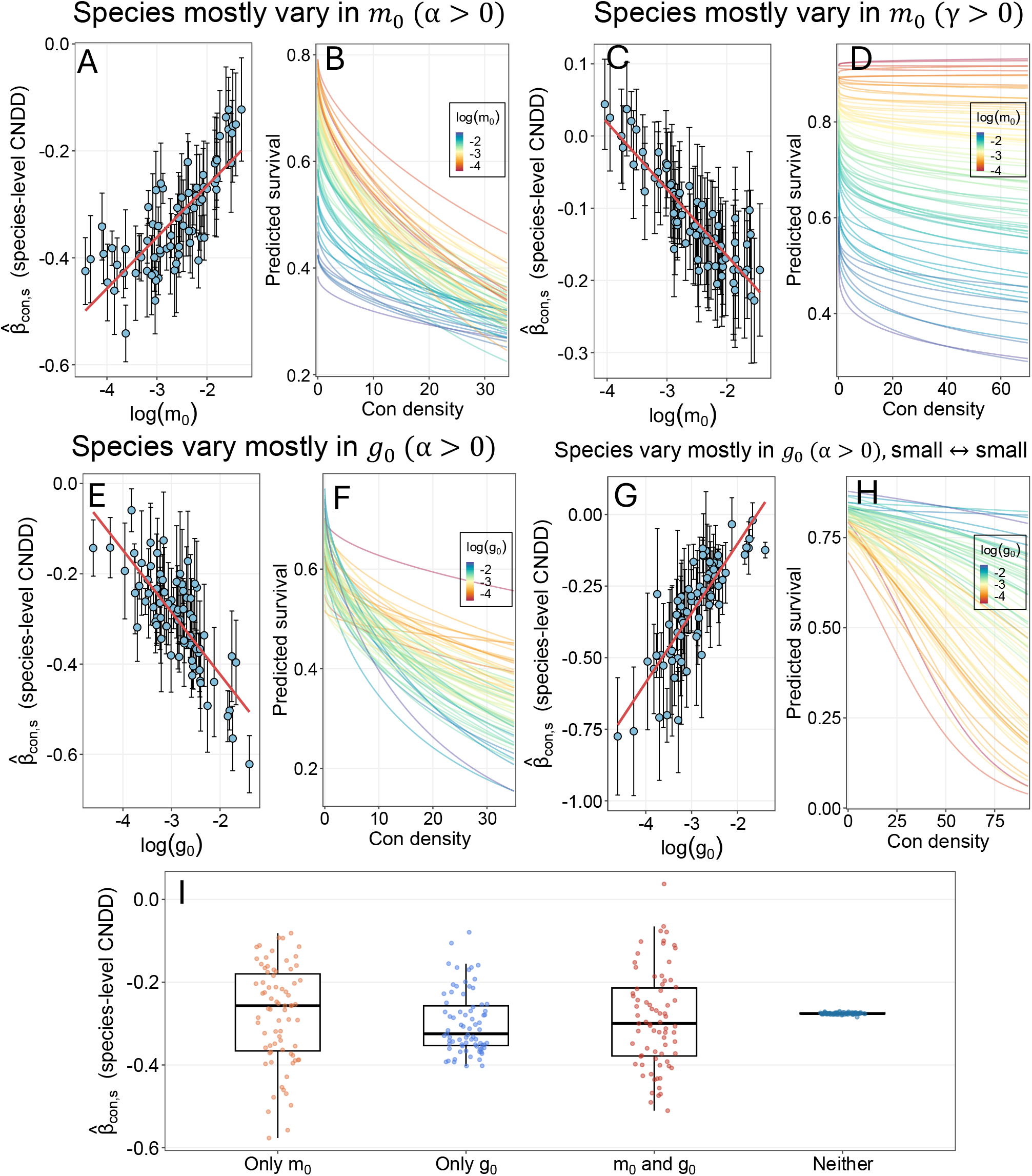
Interspecific variation in density-independent vital rates generates systematic interspecific variation in measured CNDD. GLMM fits (Eq. 9) from census survival data generated from multispecies stochastic simulations. In each panel pair (A-H), species differ primarily in the labeled baseline vital rate; variation in instantaneous density sensitivity (*α* or *γ*) is present but modest relative to the focal vital-rate variation (see Table S3). (A,C,E,G) plot the species-level conspecific density slope 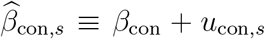 (the sum of the fixed effect and the species-specific random deviation) against the log of the focal vital-rate parameter; the red line shows the least-squares regression. (B,D,F,H) show the corresponding GLMM-implied survival–conspecific-density relationships for each species, colored by the species’ log vital-rate value. (A,B) Akin to Case 1: species differ mainly in baseline mortality rate *m*_0_; density acts through mortality (*α >* 0, *γ* = 0). (C,D) Akin to Case 2: species differ mainly in *m*_0_; density acts through growth suppression (*γ >* 0, *α* = 0). (E,F) Akin to Case 3: species differ mainly in baseline growth rate *g*_0_; density acts through mortality (*α >* 0, *γ* = 0), with density feedback dominated by larger individuals. (G,H) Akin to Case 4: species differ mainly in *g*_0_; density acts through mortality (*α >* 0, *γ* = 0), with density feedback concentrated among small individuals. (I) Variance decomposition: distributions of species-level 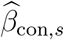 under four simulation conditions in which only *m*_0_ varies, only *g*_0_ varies, both vary, or neither varies across species, with instantaneous density sensitivity (*α >* 0) held constant across all species. Simulations contain 75 species and included data generated from 800 patches on a 10,000 patch landscape. Full details are provided in the Supplementary Information section 10; see Table S3 for all simulation parameters.

## Discussion

CNDD shapes plant diversity through its cumulative effects on juvenile survival. We show that density-independent vital rates — baseline mortality and growth — strongly modulate the strength of these cumulative effects, with the direction and magnitude depending on whether density acts through mortality or growth and which size classes are involved. These effects hold for both cohorts newly established by a reproductive pulse (e.g., for first-year seedling survival data; Lebrija-Trejos et al., 2023) and for standing populations near a dynamic equilibrium (e.g., datasets composed of larger, more established seedlings; Comita et al., 2023) — which correspond to our closed-cohort and equilibrium model variants, respectively. Consequently, CNDD variation across environments and among species systematically reflects demographic context, not just instantaneous density sensitivity. This demographic modulation of CNDD is not a statistical artifact: it reflects genuine biological differences in how conspecific crowding shapes survival across biologically relevant timescales of juvenile development.

### CNDD strength and abiotic environmental conditions

CNDD varies along abiotic environmental gradients, and such variation is typically attributed to abiotic conditions directly altering instantaneous density sensitivity. However, our results show that abiotic gradients could alternatively generate variation in CNDD by altering density-independent vital rates. The same gradient may strengthen or weaken CNDD depending on which vital rate it predominantly affects and how size affects density dependence.

Lebrija-Trejos et al. (2023) observed stronger seedling CNDD under moister conditions — a pattern interpreted as increased pathogen transmissibility in wet soils (e.g., Milici, 2024). Our results suggest an alternative hypothesis: moister soils may reduce baseline mortality and/or increase baseline growth, strengthening CNDD in this case through demographic modulation. This is illustrated by our SI pathogen model: lower baseline mortality strengthens CNDD by increasing the opportunity for infection to establish and spread, even with constant pathogen transmission (*β*) and virulence (*m*_inf_). Consistent with this interpretation, Lebrija-Trejos et al. (2023) report that soil moisture increases survival at low conspecific densities but decreases it at high densities. Figs. 4B, 4F, and S3 also show this pattern.

Similar patterns appear elsewhere: Xu et al. (2022) also found that soil moisture increases both baseline seedling survival and CNDD strength; LaManna et al. (2017) reported stronger CNDD in wet, resource-rich soils alongside higher density-independent survival; and Krishnadas and Stump (2021) found that CNDD strength and baseline seed-to-seedling survival increase with distance from the forest edge. These studies are consistent with the demographic modulation hypothesis. Similar logic applies to other gradients on which CNDD varies, like elevation (Fibich et al., 2021; LaManna et al., 2022). Demographic modulation may also help explain why measured CNDD effects change after conspecific adult mortality (‘legacy CNDD’; e.g., Magee et al., 2025): the post-mortality environment may alter density-independent vital rates, thereby influencing realized interval CNDD.

Crucially, environmental variation in measured CNDD could reflect heterogeneity in (i) instantaneous density sensitivity (*α* or *γ*) or (ii) demographic modulation via density-independent vital rates. A simple diagnostic experiment could test whether the latter is feasible: apply an exogenous, density-independent mortality hazard by removing seedlings at random at rate *m* throughout the census interval, independent of local conspecific density, with *m* varied across plots. A significant *m×*conspecific density interaction in a GLMM fit to the resulting survival data would demonstrate that variation in density-independent mortality is sufficient to generate variation in CNDD.

### Inter-specific variation in CNDD and life history trade-offs

Interspecific variation in measured CNDD is well-documented and typically interpreted as reflecting factors such as differences among species in their susceptibility to natural enemies or intraspecific competition (e.g., LaManna et al., 2016; Stump and Comita, 2018). Our results show that interspecific differences in density-independent vital rates generate structured variation in CNDD. However, the same vital rate may affect CNDD in opposite ways depending on (i) how size affects density-dependent effects and susceptibility, and (ii) which vital rate density influences; accurately interpreting vital rate–CNDD relationships requires understanding the underlying demographic pathway.

These patterns help contextualize measured CNDD within life-history trade-offs. Several studies indicate that faster-growing species exhibit stronger measured CNDD than slower-growing species (Lebrija-Trejos et al., 2016; Zhu et al., 2018; Song et al., 2021*b*; Zang et al., 2021; Qin et al., 2022; Huanca-Nunez et al., 2026), a relationship attributed to a defense– growth trade-off in instantaneous density sensitivity: fast-growing species invest less in defense, making them more susceptible to natural enemies (Coley, 1980; McCarthy-Neumann and Kobe, 2008). Our results show that this pattern can emerge simply because faster growth rate *causes* stronger measured CNDD (Cases 2–3; although see Case 4).

Because growth and mortality rates are themselves correlated across species (Wright et al., 2010), Zhu et al. (2018) and Huanca-Nunez et al. (2026) link stronger measured CNDD not just to faster growth but to species’ position along the growth–mortality trade-off axis (faster growth, higher mortality). Our framework offers a direct explanation for this pattern when density suppresses growth (Case 2): here, both higher baseline mortality and higher baseline growth strengthen realized interval CNDD, so species further along the trade-off axis necessarily experience stronger CNDD. When density instead acts directly on mortality (Case 1), however, higher baseline mortality weakens CNDD — the opposite of the empirical association outlined by Zhu et al. (2018) and Huanca-Nunez et al. (2026). This tension potentially resolves by recognizing that the mortality measurements in these studies (e.g., Wright et al., 2010) are raw mortality probabilities that include density-dependent components, not density-independent mortality (i.e.,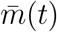). In fact, species with faster growth may experience higher measured mortality because they experience stronger CNDD, raising the possibility that the growth–mortality trade-off is partly an emergent consequence of density-dependent feedbacks. Similar logic applies for CNDD associations with other life-history traits, such as shade tolerance (Brown et al., 2020), drought tolerance (Song et al., 2024), and species position along the stature–recruitment trade-off (Huanca-Nunez et al., 2026).

### CNDD and community structure

Our results have implications for species coexistence. Recruitment to adulthood is the product of survival across successive life stages (Fig. 1F); our results show that CNDD within each stage is shaped by density-independent vital rates. This matters because theoretical papers assessing how CNDD affects coexistence typically assume CNDD is independent of fitness differences arising from density-independent vital rates (e.g., Chisholm and Muller-Landau, 2011; Stump and Chesson, 2015; Chisholm and Fung, 2020; Smith, 2022*a*,*b*; Magee et al., 2025) — an assumption our models contradict. This is particularly relevant given that interspecific variation in CNDD can erode diversity: past models show that species experiencing relatively strong CNDD are at a disadvantage (Miranda et al., 2015; Stump and Comita, 2018; May et al., 2020). However, a species may experience stronger CNDD precisely because it has lower density-independent mortality or higher density-independent growth — both of which are competitive advantages, all else equal. Interspecific CNDD variation driven by density-independent vital rates may in fact equalize interspecific fitness differences, facilitating species coexistence. Analyses linking empirical CNDD estimates to coexistence models should therefore not treat fitted coefficients (*β*_1_ or *β*_con,*s*_) as direct proxies for phenomenological density-dependence parameters. Minimally, such analyses should test whether measured CNDD covaries with independently estimated baseline vital rates. Correctly parameterizing coexistence models, however, requires estimating density-independent vital rates as functions of size and instantaneous density sensitivity (*α* or *γ*) as distinct parameters, rather than collapsing both into a single regression coefficient that entangles them.

## Conclusions

CNDD is shaped by demography in predictable ways. When CNDD varies across abiotic gradients or species, variation in density-independent vital rates should be considered alongside variation in instantaneous density sensitivity as an explanation. Distinguishing between these requires identifying the demographic pathway through which density acts rather than relying on measured CNDD alone as a proxy for per-capita sensitivity to conspecific density, a distinction with implications for how CNDD maintains species diversity.

## Acknowledgments

This research was funded by the U.S. National Science Foundation grant NSF OCE 1851489 and National Science Foundation Division of Environmental Biology Award 2240430. I thank J. Timothy Wootton, Catherine A Pfister, Mercedes Pascual, Trevor Price, Lukas Magee, and the Wootton-Pfister lab for insightful discussion and feedback on the manuscript.

## Authorship

D.J.B.S. conceived the study, conducted the analyses, and wrote the first draft of the manuscript. D.J.B.S and A.O. edited revisions. D.J.B.S, B.E.S., D.F., and A.O. contributed to conceptual development and revisions.

## Data accessibility statement

All code and data supporting the results are archived on figshare and available at: https://doi.org/10.6084/m9.figshare.33055142

