## Supplementary Information for "What drives variation in conspecific negative density-dependence? A demographic perspective"

### Supplementary Information: What generates variation in conspecific negative density-dependence?

Daniel J. B. Smith*
The Oden Institute, The University of Texas at Austin, Austin, TX USA

5.4 Case 3: Large individuals drive density-dependent mortality of small individuals 23
5.5 Case 4: Small individuals generate and experience density-dependent mortality 25

**6 Instantaneous density sensitivity and realized interval CNDD** **27**

**7 Case 2: the effect of baseline growth $g_0$** **29**

**8 Equilibrium analysis of PDE model** **30**

**11 GAM robustness analysis** **41**

**12 Simulation parameters for Fig. 3, Fig. S2, and Fig. S4** **44**

**13 Simulation parameters for Figures 4 and 5** **46**

**14 Variable / parameter table** **49**

### Overview

This supplement provides derivations, numerical validation, and parameter details supporting the results presented in the main text. Section 1 derives the general relationship between cumulative hazard and the logit-scale regression slope $\beta_1$ used throughout the main text and Box 1. Sections 2–4 treat the SI pathogen model: deriving the infection establishment probability quoted in Results, the closed-form approximation for $\beta_1(m)$, and the model's endemic equilibrium behavior. Section 5 derives the four two-stage approximations reported in Box 1, and Section 6 confirms numerically that greater instantaneous density sensitivity steepens realized interval CNDD, independently of the baseline-rate effects that are the main text's primary focus. Section 7 numerically confirms the effect of baseline growth $g_0$ on CNDD in Case 2. Section 8 extends the size-structured model's closed-cohort results to standing, equilibrium populations. Section 9 shows that the Gillespie SSA used in the spatially explicit multispecies simulations recovers a finite-volume approximation of the size-structured PDE in its mean-field limit. Section 10 details the spatially explicit multispecies simulations and GLMM fitting procedure. Section 11 presents a GAM-based robustness check of the interspecific CNDD patterns in Fig. 5. Parameter tables and full simulation details for all main-text and Supplementary figures are provided in Sections 12–13, and Section 14 provides a full glossary of notation used throughout the main text and Supplement.

### 1 From cumulative hazard to the empirical regression slope

We derive the connection between cumulative hazard and the logit-scale regression slope $\beta_1$ used throughout the main text and Box 1. This relationship underlies its application to the SI pathogen model below (Section 3) and the derivations related to the two-stage approximation of the Partial Differential Equation (PDE) model in Section 5.

Empirical studies commonly model survival probability $p_i$ of individual $i$ as $\text{logit}(p_i) = \beta_0 + \beta_1 N_i(0) + \ldots$, with $\beta_1 < 0$ indicating CNDD. Since cohort survival is $p_i = \exp(-\text{H}(T))$, we have

$$\text{logit}(p_i) = \log\left(\frac{p_i}{1-p_i}\right) = \log\left(\frac{e^{-\text{H}(T)}}{1-e^{-\text{H}(T)}}\right) = -\text{H}(T) - \log\bigl(1 - e^{-\text{H}(T)}\bigr)\,.$$

Differentiating with respect to $N(0)$,

$$\frac{\partial}{\partial N(0)}\text{logit}(p_i) = -\frac{\partial \text{H}(T)}{\partial N(0)}\left[1 + \frac{e^{-\text{H}(T)}}{1-e^{-\text{H}(T)}}\right] = -\frac{\partial \text{H}(T)}{\partial N(0)} \cdot \frac{1}{1-e^{-\text{H}(T)}}.$$

Multiplying numerator and denominator by $e^{\text{H}(T)}$,

$$\beta_1 \approx \frac{\partial}{\partial N(0)}\text{logit}(p_i) = -\frac{e^{\text{H}(T)}}{e^{\text{H}(T)}-1} \cdot \frac{\partial \text{H}(T)}{\partial N(0)}. \tag{S1}$$

This is exact when logit-survival is linear in $N(0)$, and otherwise approximates the local logit-scale density effect, which an estimated $\hat{\beta}_1$ recovers as a weighted average over the observed density range.

**High-survival simplification.** When $\mathrm{H}(T) \to 0$ (survival near 1), $e^{\mathrm{H}(T)} \approx 1 + \mathrm{H}(T)$, so $\frac{e^{\mathrm{H}(T)}}{e^{\mathrm{H}(T)}-1} \approx \frac{1}{\mathrm{H}(T)}$, and Eq. S1 reduces to

$$\beta_1 \approx -\frac{1}{\mathrm{H}(T)} \frac{\partial \mathrm{H}(T)}{\partial N(0)}. \tag{S2}$$

This is the approximation used in the SI pathogen model below (Section 3) and throughout Box 1 and the four two-stage cases in Section 5: each application derives $\partial \mathrm{H}(T)/\partial N(0)$ and $\mathrm{H}(T)$ at leading order in $T$, then substitutes both into Eq. S2 to obtain $\beta_1$ in terms of model parameters.

### 2 Establishment probability in the SI pathogen model

We derive the analytical approximation for infection establishment probability presented in the main text for the Susceptible Infected (SI) model. We restate the model for completeness, then derive the basic and effective reproduction numbers, the Poisson thinning argument for establishment probability, and the closed-form and near-threshold expressions quoted in Results.

#### 2.1 Model and goal

The following recaps the SI model introduced in the main text. We consider a continuous-time stochastic susceptible–infected (SI) model of pathogen dynamics within a seedling cohort. Each simulation begins with a cohort of $N(0)$ seedlings, all susceptible ($S(0) = N(0)$, $I(0) = 0$). Susceptible individuals become infected either via background exposure at rate $\xi S$ or via mass-action transmission from infected individuals at rate $\beta SI$. Both susceptible and infected individuals die at baseline rate $m$; infected individuals additionally die at rate $m_{\mathrm{inf}}$. The mean-field equations are

$$\frac{dS}{dt} = f - \xi S - \beta SI - mS, \qquad \frac{dI}{dt} = \xi S + \beta SI - (m + m_{\mathrm{inf}})I \tag{S3}$$

where $f$ is seedling supply rate. In this analysis, we assume the closed-cohort model, so we set $f = 0$. In the main text we show that measured CNDD — the fitted logit-scale density slope $\hat{\beta}_1(m)$ from interval-census GLMs — weakens as baseline mortality $m$ increases, and we attribute this to suppression of infection establishment. Here we derive an analytical approximation for the probability that infection establishes within a census interval $[0, T]$, $P_{\mathrm{est}}(T)$, and show explicitly how it depends on $m$ and $N(0)$.

#### 2.2 The basic and effective reproduction numbers

The key quantity governing whether an infection can spread from a single infected individual is the reproduction number — the expected number of secondary infections it produces before being removed.

At the very start of the interval, the full cohort is susceptible and $S(0) = N(0)$. A single infected individual produces new infections at rate $\beta S(0) = \beta N(0)$ and is removed (by death) at rate $m + m_{\mathrm{inf}}$. Their ratio defines the *basic reproduction number*,

$$R_0 \equiv \frac{\beta N(0)}{m + m_{\mathrm{inf}}}, \tag{S4}$$

which measures the initial potential for spread and depends explicitly on both initial cohort size $N(0)$ and baseline mortality $m$. When $R_0 > 1$ an introduced infection deterministically spreads; when $R_0 \leq 1$ it does not.

As time progresses, susceptibles are lost to background mortality before any outbreak occurs, so the potential for spread declines. Approximating $S(t) \approx N(0)e^{-mt}$ (valid before any large outbreak), a single infected individual introduced at time $t$ produces secondary infections at rate $\beta S(t)$ and is removed at rate $m + m_{\mathrm{inf}}$. The *effective reproduction number at time* $t$ is therefore

$$R_t \equiv \frac{\beta S(t)}{m + m_{\mathrm{inf}}} \approx \frac{\beta N(0)}{m + m_{\mathrm{inf}}}\, e^{-mt} = R_0\, e^{-mt}. \tag{S5}$$

Thus $R_t$ is simply $R_0$ reduced by the fraction of the cohort that remains susceptible at time $t$. Because $R_t$ is strictly decreasing in $t$, there exists a finite *critical time*

$$t_c = \frac{1}{m} \log R_0, \tag{S6}$$

beyond which $R_t < 1$ and no introduced infection can spread. For $t > t_c$, establishment is impossible regardless of transmission parameters.

#### 2.3 Establishment probability: Poisson thinning argument

We approximate the process of successful infection establishments as a thinned Poisson process. Background introductions (single infected individuals appearing from external sources) arrive at rate

$$\lambda_{\text{intro}}(t) = \xi S(t) \approx \xi N(0)\, e^{-mt}. \tag{S7}$$

Conditional on an introduction at time $t$, we approximate early spread by a linear birth–death branching process with per-capita birth rate $b(t) = \beta S(t)$ and death rate $d = m + m_{\text{inf}}$. For such a process initiated by one individual, the probability of non-extinction (establishment) is

$$q(t) = \Pr(\text{establish} \mid \text{introduced at } t) \approx \max\left\{0,\ 1 - \frac{1}{R_t}\right\}. \tag{S8}$$

the classical extinction-probability result for linear birth–death processes (Allen, 2008, Eqs. 3.15–3.16). This approximates $q(t)$ by its value under the constant-rate (time-homogeneous) case, treating $S(t)$ as effectively frozen at its value at the moment of introduction — valid when whether an introduced infection takes off is decided quickly relative to the timescale of susceptible depletion, $1/m$ (i.e., pathogen spread is fast relative to mean density-independent seedling mortality). Fig. 2E validates this approximation directly against simulated establishment probabilities.

Since $q(t) = 0$ for $t \geq t_c$, only introductions before $\tau = \min(T, t_c)$ can establish. Treating successful establishments as a thinned Poisson process with intensity $r(t) = \lambda_{\text{intro}}(t)\, q(t)$, the probability of at least one establishment by time $T$

$$P_{\text{est}}(T) \approx 1 - \exp\left[-\int_0^{\tau} r(t)\, dt\right] \tag{S9}$$

recalling $\tau = \min(T, t_c)$

#### 2.4 Closed-form expression

For $t \le t_c$, using $R_t = R_0 e^{-mt}$,

$$r(t) = \xi N(0)\, e^{-mt} \left(1 - \frac{e^{mt}}{R_0}\right) = \xi N(0) \left(e^{-mt} - \frac{1}{R_0}\right). \tag{S10}$$

Integrating from 0 to $\tau$,

$$\int_0^{\tau} r(t)\, dt = \xi N(0) \left(\frac{1 - e^{-m\tau}}{m} - \frac{\tau}{R_0}\right), \tag{S11}$$

giving the closed-form approximation

$$P_{\text{est}}(T) \approx 1 - \exp\left[-\xi N(0) \left(\frac{1 - e^{-m\tau}}{m} - \frac{\tau}{R_0}\right)\right], \qquad \tau = \min(T,\, t_c). \tag{S12}$$

Eq. S12 makes explicit the two routes by which higher $m$ suppresses establishment: it reduces introduction opportunities by depleting susceptibles faster ($\lambda_{\text{intro}}(t) \propto e^{-mt}$), and it shrinks the supercritical window by reducing $t_c = (1/m) \log R_0$.

#### 2.5 Near-threshold simplification

When $T \ge t_c$ (the full supercritical window falls within the census interval) and $R_0$ is close to its threshold value of 1, a further simplification is possible. Let

$$x \equiv 1 - \frac{1}{R_0}, \qquad 0 < x \ll 1 \text{ near threshold}, \tag{S13}$$

so that $R_0 = 1/(1-x)$. Using the Mercator series $-\log(1-x) = x + x^2/2 + O(x^3)$,

$$\log R_0 = -\log(1-x) = x + \frac{x^2}{2} + O(x^3), \tag{S14}$$

so that

$$\frac{\log R_0}{R_0} = (1-x)\left(x + \frac{x^2}{2} + O(x^3)\right) = x - \frac{x^2}{2} + O(x^3). \tag{S15}$$

Since $1 - 1/R_0 = x$ by definition, subtracting gives

$$\left(1-\frac{1}{R_0}\right) - \frac{\log R_0}{R_0} = x - \left(x - \frac{x^2}{2}\right) + O(x^3) = \frac{1}{2}\left(1-\frac{1}{R_0}\right)^2 + O\left(\left(1-\frac{1}{R_0}\right)^3\right), \tag{S16}$$

so that the integral over $[0, t_c]$ reduces to

$$\int_0^{t_c} r(t)\, dt = \frac{\xi N(0)}{m}\left[\left(1-\frac{1}{R_0}\right) - \frac{\log R_0}{R_0}\right] \approx \frac{\xi N(0)}{2m}\left(1-\frac{1}{R_0}\right)^2. \tag{S17}$$

This yields the near-threshold approximation quoted in the main text,

$$P_{\text{est}}(T) \approx 1 - \exp\left[-\frac{\xi N(0)}{2m}\left(1-\frac{1}{R_0}\right)^2\right], \qquad (T \geq t_c,\ R_0 \approx 1), \tag{S18}$$

in which the dependence on $m$ in the exponent is transparent: holding $R_0$ fixed, $P_{\text{est}}(T)$ decreases monotonically with $m$.

### 3 Approximation for the regression slope $\beta_1(m)$ in the SI model

We derive the closed-form approximation for the logit-scale density slope $\beta_1(m)$ quoted in the main text Results, using the general hazard-to-regression-slope relationship derived in Section 1 above (Eq. S2). To do so, we derive approximations for $H(T)$ and $\partial \mathrm{H}(T)/\partial N(0)$ for the closed-cohort Susceptible-Infected (SI) model analyzed in the main text.

#### 3.1 Small-time expansion of the infection hazard

We use the closed-cohort SI model (Eq. S3) with $S(0) = N(0)$, $I(0) = 0$. The instantaneous hazard is $h(t) = m + m_{\text{inf}}\, I(t)/N(t)$ (main text Eq. 11), so the cumulative hazard decomposes as

$$\mathrm{H}(T) = mT + m_{\text{inf}} \int_0^T \frac{I(t)}{N(t)}\, dt.$$

Since $I(0) = 0$, the ratio $I(t)/N(t)$ must build up from zero; we obtain its short-time behavior by Taylor-expanding $I(t)$ and $N(t)$ about $t = 0$.

From $\dot{I} = \xi S + \beta S I - (m + m_{\text{inf}})I$ and $I(0) = 0$,

$$\dot{I}(0) = \xi N(0), \qquad \ddot{I}(0) = \xi N(0)\left(\beta N(0) - \xi - 2m - m_{\text{inf}}\right),$$

so

$$I(t) = \xi N(0)\, t + \frac{1}{2}\xi N(0)\left(\beta N(0) - \xi - 2m - m_{\text{inf}}\right) t^2 + O(t^3).$$

Similarly, from $\dot{N} = -mN - m_{\text{inf}} I$ and $N(0) = N(0)$,

$$N(t) = N(0)\,(1 - mt) + O(t^2).$$

Dividing, the baseline mortality term cancels at this order:

$$\frac{I(t)}{N(t)} = \xi t + \frac{1}{2}\xi\left(\beta N(0) - \xi - m_{\text{inf}}\right)t^2 + O(t^3). \tag{S19}$$

#### 3.2 Cumulative hazard and its sensitivity to $N(0)$

Integrating Eq. S19 from 0 to $T$ and adding the baseline term,

$$\text{H}(T) = mT + \frac{1}{2}m_{\text{inf}}\xi T^2 + \frac{1}{6}m_{\text{inf}}\xi\left(\beta N(0) - \xi - m_{\text{inf}}\right)T^3 + O(T^4). \tag{S20}$$

The density-independent part of the cumulative hazard, denoted $\text{H}_{\text{DI}}(T)$, is obtained by dropping the $N(0)$-dependent term:

$$\text{H}_{\text{DI}}(T) \approx mT + \frac{1}{2}m_{\text{inf}}\xi T^2. \tag{S21}$$

Differentiating Eq. S20 with respect to $N(0)$ gives the leading-order sensitivity,

$$\frac{\partial \text{H}(T)}{\partial N(0)} \approx \frac{1}{6}\beta m_{\text{inf}}\xi T^3. \tag{S22}$$

As in Cases 3–4 of Section 5, there is no $O(T)$ or $O(T^2)$ term: because $I(0) = 0$, the density-dependent contribution to the hazard must first be seeded via $\xi$ before mass-action transmission can act, delaying the onset of the density signal to $O(T^3)$.

#### 3.3 Regression coefficient

Substituting Eqs. S21 and S22 into the general relationship $\beta_1 \approx -\left[\partial\text{H}(T)/\partial N(0)\right]/\text{H}(T)$ (Eq. S2), and using $\text{H}_{\text{DI}}(T)$ in place of the full $\text{H}(T)$ in the denominator (valid since the $N(0)$-dependent term in $\text{H}(T)$ is already higher-order), gives

$$\beta_1(m) \approx -\frac{\beta\, m_{\text{inf}}\,\xi\, T^2}{6m + 3m_{\text{inf}}\,\xi\, T}. \tag{S23}$$

This is the approximation quoted in the main text (Eq. 15).

### 4 Equilibrium analysis of the SI model with seedling supply

We derive the endemic equilibrium of the SI model with seedling supply and show that baseline mortality $m$ shapes the per-capita density-dependent hazard through equilibrium infection prevalence, independently of any effect on total equilibrium density. Results are illustrated in Fig. S1.

#### 4.1 Model

For analytical tractability, we set $\xi = 0$ and consider the simplified supply model

$$\frac{dS}{dt} = f - \beta SI - mS, \qquad \frac{dI}{dt} = \beta SI - (m + m_{\text{inf}})I, \tag{S24}$$

where $f$ is a constant seedling supply rate, $\beta$ is the mass-action transmission rate, $m$ is the baseline per-capita mortality rate, and $m_{\text{inf}}$ is the additional mortality of infected individuals. Stochastic simulations in Fig. S1E–F retain $\xi > 0$ as in the main text.

#### 4.2 Endemic equilibrium

Setting $dS/dt = dI/dt = 0$ in Eq. S24 and solving, we obtain the endemic equilibrium

$$S^* = \frac{m + m_{\text{inf}}}{\beta}, \qquad I^* = \frac{f}{m + m_{\text{inf}}} - \frac{m}{\beta}, \qquad N^* = S^* + I^* = \frac{m_{\text{inf}}}{\beta} + \frac{f}{m + m_{\text{inf}}}. \tag{S25}$$

The endemic equilibrium exists (i.e., $I^* > 0$) when $\beta f > m(m + m_{\text{inf}})$, which is the condition that the pathogen can invade a fully susceptible population supplied at rate $f$.

##### 4.3 Per-capita density-dependent hazard at equilibrium

At endemic equilibrium, the excess per-capita mortality hazard due to infection experienced by a randomly chosen focal individual is

$$m_{\mathrm{DD}}^{*} \equiv \frac{m_{\mathrm{inf}} \cdot I^{*}}{N^{*}} = m_{\mathrm{inf}} \cdot \frac{I^{*}}{N^{*}}, \tag{S26}$$

which is the infection fatality rate weighted by equilibrium prevalence $I^{*}/N^{*}$. This quantity is directly analogous to $\bar{m}_{\mathrm{DD}}(t)$ in the cohort analysis of the main text, and is shown as a function of $N^{*}$ for varying $m$ in Fig. S1A–B. Substituting Eq. S25 into Eq. S26 yields

$$m_{\mathrm{DD}}^{*} = \frac{m_{\mathrm{inf}}(\beta f - m(m + m_{\mathrm{inf}}))}{m_{\mathrm{inf}}(m + m_{\mathrm{inf}}) + \beta f}. \tag{S27}$$

Crucially, smaller $m$ makes $m_{\mathrm{DD}}^{*}$ increase more rapidly with equilibrium density $N^{*}$ (Fig. S1A-B).

##### 4.4 Survival consequences

The density-dependent component of interval survival is $\exp(-m_{\mathrm{DD}}^{*})$ per unit time, shown in Fig. S1C as a function of $N^{*}$: lower $m$ produces a lower survival probability at any given equilibrium density, with the separation between $m$ values widening as $N^{*}$ increases. Total survival over the census interval $[0, T]$ — including both baseline and density-dependent mortality — is approximately $\exp(-(m + m_{\mathrm{DD}}^{*})\,T)$, shown on a log scale in Fig. S1D. Higher baseline mortality $m$ reduces total survival directly, but also reduces $m_{\mathrm{DD}}^{*}$, partially offsetting its own effect on total survival; the net result is that curves for different $m$ are more compressed on the log scale than the density-dependent component alone would suggest.

#### 4.5 Effect of baseline mortality

**Total effect.** Differentiating Eq. S27 with respect to $m$ using the quotient rule gives

$$\frac{\partial m_{\mathrm{DD}}^*}{\partial m} = \frac{-m_{\mathrm{inf}}(m + m_{\mathrm{inf}})(m_{\mathrm{inf}}(m + m_{\mathrm{inf}}) + 2\beta f)}{(m_{\mathrm{inf}}(m + m_{\mathrm{inf}}) + \beta f)^2} < 0, \tag{S28}$$

which is strictly negative whenever the endemic equilibrium exists. Higher baseline mortality therefore always reduces the equilibrium density-dependent hazard (Fig. S1A–B).

**Effect holding $N^*$ fixed.** To isolate the prevalence pathway, we hold $N^*$ fixed. Since $S^* = (m + m_{\mathrm{inf}})/\beta$ does not depend on $f$,

$$m_{\mathrm{DD}}^*\big|_{N^*\text{ fixed}} = m_{\mathrm{inf}}\left(1 - \frac{m + m_{\mathrm{inf}}}{\beta N^*}\right), \tag{S29}$$

and differentiating with respect to $m$,

$$\left.\frac{\partial m_{\mathrm{DD}}^*}{\partial m}\right|_{N^*\text{ fixed}} = -\frac{m_{\mathrm{inf}}}{\beta N^*} < 0. \tag{S30}$$

This is unconditionally negative: higher baseline mortality increases the removal rate of infected individuals, requiring a higher equilibrium susceptible density $S^*$ to sustain transmission. Holding $N^*$ fixed, this leaves a smaller infected fraction at equilibrium (Fig. S1E), directly reducing $m_{\mathrm{DD}}^*$.

#### 4.6 Simulation procedure

To confirm that the analytical results extend to stochastic dynamics, we generated census-style survival data from the stochastic version of Eq. S24, simulated using a Gillespie SSA with $\xi > 0$ as in the main text. For each value of baseline mortality $m$, we selected the seedling supply rate $f$ to target a grid of equilibrium densities $N^*$ (using the approximation $N^* \approx m_{\mathrm{inf}}/\beta + f/(m + m_{\mathrm{inf}})$ from Eq. S25 to solve for $f$). For each $(m, N^*)$ combination,

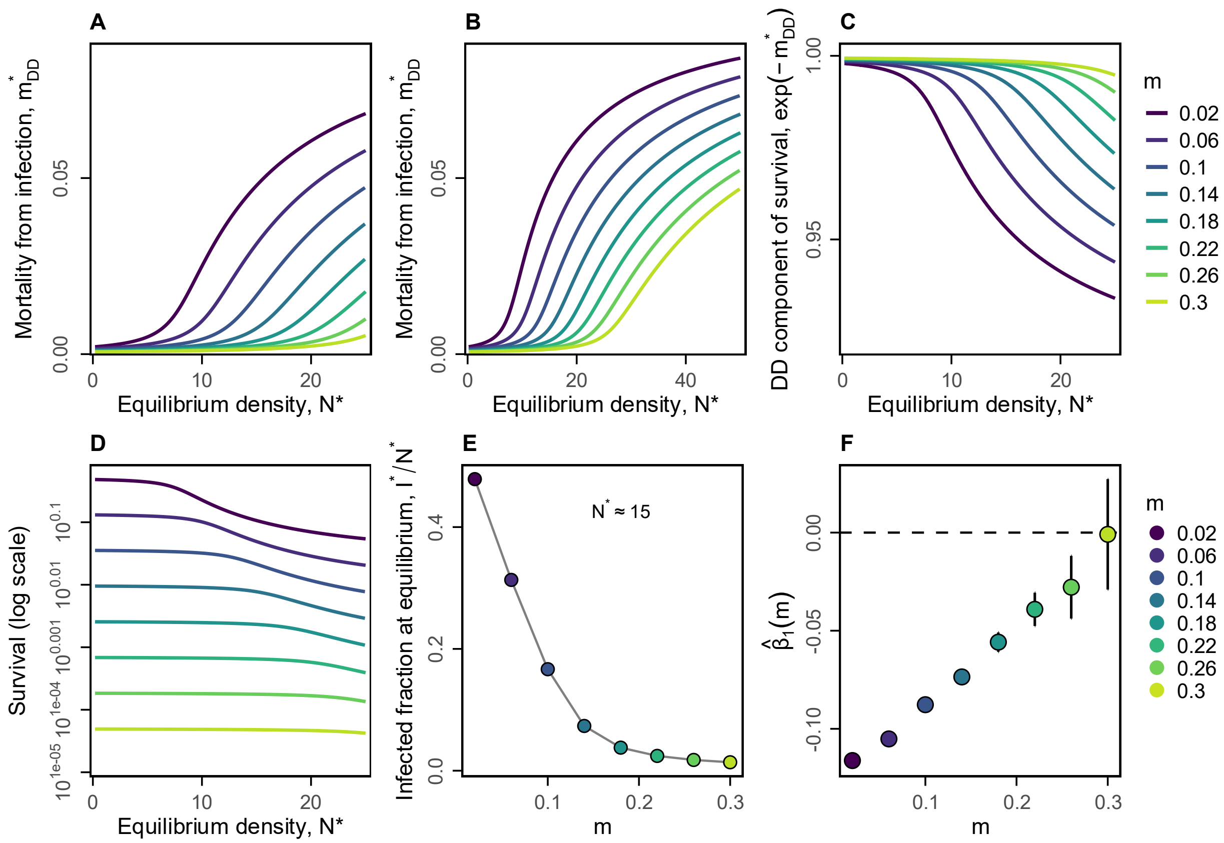

Figure S1: **Equilibrium SI model: baseline mortality weakens pathogen-mediated CNDD through infection prevalence.** (A) Excess mortality rate due to infection, $m^*_{\mathrm{DD}} = m_{\mathrm{inf}} I^*/N^*$, as a function of equilibrium density $N^*$ for each value of baseline mortality $m$ (colors; see legend), shown over a restricted density range. (B) Same as (A) over a wider density range. (C) Density-dependent component of per-unit-time survival, $\exp(-m^*_{\mathrm{DD}})$, as a function of $N^*$. (D) Total interval survival, $\exp(-(m+m^*_{\mathrm{DD}})\,T)$, on a log scale, as a function of $N^*$. (E) Equilibrium infected fraction $I^*/N^*$ as a function of $m$, evaluated at fixed equilibrium density $N^* \approx 15$ (by choosing $f$ to target this density for each $m$). (F) Measured CNDD ($\hat{\beta}_1$) from GLMs fitted to census data generated from stochastic equilibrium simulations, as a function of $m$. Points show estimates $\pm 1.96\,$SE. CNDD weakens monotonically as $m$ increases, mirroring the cohort result in Fig. 2. Parameters: $\xi = 0.0025$, $\beta = 0.015$, $m_{\mathrm{inf}} = 0.1$, $T = 33$.

we ran a Gillespie burn-in of duration $T_{\text{burn}} = 50$ time units to allow the system to approach steady state, then tracked the fate of the cohort present at the end of the burn-in over a census interval of length $T = 33$. To separate original cohort members from new arrivals, we tracked four compartments: susceptible and infected individuals present at the start of the census interval ($S_{\text{orig}}$, $I_{\text{orig}}$), and susceptible and infected new arrivals ($S_{\text{new}}$, $I_{\text{new}}$). New arrivals interacted with but were not counted in the cohort. We ran 10000 independent replicates per $(m, N^*)$ cell and fitted the same binomial GLM as in the main text to the resulting survival outcomes, yielding $\hat{\beta}_1$ as a function of $m$ (Fig. S1F). All parameters match those used in the main text ($\xi = 0.0025$, $\beta = 0.015$, $m_{\text{inf}} = 0.1$).

Consistent with the analytical results, fitting GLMs to census data generated from equilibrium simulations yields $\hat{\beta}_1$ values that become progressively less negative as $m$ increases (Fig. S1F), mirroring the cohort result in the main text.

### 5 Two-stage approximations and $\beta_1$ derivations (Box 1)

We derive the approximations reported in Box 1 of the main text. We collapse the continuous size-structured model (Eq. 4 of the main text) to a minimal two-stage structure in order to isolate, analytically, how baseline demography and the demographic channel through which density acts shape realized interval CNDD. The four cases correspond directly to the four cases in Box 1, and the results are reproduced here with full derivation steps, resulting in the corresponding $\beta_1$ expression obtained via Eq. S2.

#### 5.1 Setup and notation

We track a cohort of juveniles in two stages, "small" ($J_1$) and "large" ($J_2$), with per-capita transition rate $g(J_1, J_2)$ from small to large and stage-specific per-capita mortality hazards $\mu_1(J_1, J_2)$ and $\mu_2(J_1, J_2)$:

$$\frac{dJ_1}{dt} = -g(J_1, J_2)\, J_1 - \mu_1(J_1, J_2)\, J_1, \tag{S31}$$

$$\frac{dJ_2}{dt} = \quad g(J_1, J_2)\, J_1 - \mu_2(J_1, J_2)\, J_2, \tag{S32}$$

with $J_1(0) = N(0)$ and $J_2(0) = 0$. Total cohort abundance is $N(t) = J_1(t) + J_2(t)$. Adding Eqs. S31–S32,

$$\frac{dN}{dt} = -\bar{m}(t)\, N(t), \qquad \bar{m}(t) = \frac{\mu_1 J_1 + \mu_2 J_2}{J_1 + J_2}, \tag{S33}$$

where $\bar{m}(t)$ is the abundance-weighted mean hazard. Integrating,

$$\mathrm{H}(T) \equiv -\log\left(\frac{N(T)}{N(0)}\right) = \int_0^T \bar{m}(t)\, dt. \tag{S34}$$

A central quantity is the sensitivity of cumulative survival to initial cohort size,

$$\frac{\partial \mathrm{H}(T)}{\partial N(0)}, \tag{S35}$$

which we substitute, together with $\mathrm{H}(T)$ at leading order in $T$, into Eq. S2 to obtain $\beta_1$ for each case. Note that when $\alpha = \gamma = 0$, cohort survival does not depend on $N(0)$, so $\frac{\partial}{\partial N(0)}\mathrm{H}(T) = 0$ in the absence of density dependence. Our sensitivity measure therefore captures only the density-dependent contribution to cumulative hazard.

**Small-$T$ expansions.** In Cases 2–4 the density-dependent contribution to $\bar{m}$ is absent or zero at $t = 0$ and must build up over time, so we use Taylor expansions in $T$ to identify leading-order behavior. Throughout, $T \ll 1$ means $T$ is small relative to the characteristic demographic timescales set by baseline hazards and transition rates, so $J_2(t) = O(t)$ and $J_1(t) = N(0) + O(t)$ initially.

#### 5.2 Case 1: Symmetric density-dependent mortality, density-independent transition

**Assumptions.**

$$g(J_1, J_2) = g_0, \qquad \mu_1 = \mu_2 = m + \alpha N, \qquad N = J_1 + J_2.$$

Because both stages share the same hazard, stage structure is irrelevant and $N(t)$ satisfies

$$\frac{dN}{dt} = -(m + \alpha N)\, N. \tag{S36}$$

**Exact solution.** This is a Bernoulli equation; substituting $u = 1/N$ gives the linear ODE $\dot{u} = mu + \alpha$, with solution

$$N(t) = \frac{N(0)\, e^{-mt}}{1 + \dfrac{\alpha N(0)}{m}\,(1 - e^{-mt})}. \tag{S37}$$

**Cumulative hazard and sensitivity.** Using $\mathrm{H}(T) = -\log[N(T)/N(0)]$,

$$\mathrm{H}(T) = mT + \log\left(1 + \frac{\alpha N(0)}{m}\left(1 - e^{-mT}\right)\right).$$

Differentiating with respect to $N(0)$ gives the exact sensitivity,

$$\frac{\partial \mathrm{H}(T)}{\partial N(0)} = \frac{\alpha(1 - e^{-mT})}{m + \alpha N(0)(1 - e^{-mT})}, \tag{S38}$$

which is positive for all $N(0) \geq 0$.

**Small-$T$ expansion.** Substituting $1 - e^{-mT} = mT - \frac{m^2T^2}{2} + O(T^3)$ into Eq. (S38),

$$\frac{\partial \mathrm{H}(T)}{\partial N(0)} = \alpha T - \left(\frac{\alpha m}{2} + \alpha^2 N(0)\right) T^2 + O(T^3). \tag{S39}$$

The $O(T)$ term depends only on $\alpha$; baseline mortality $m$ enters at $O(T^2)$, weakening the density signal as it increases. The transition rate $g_0$ does not appear because stages are demographically identical.

**Regression coefficient.** At leading order, $\mathrm{H}(T) \approx (m + \alpha N(0))T$. Substituting this and Eq. (S39) into Eq. (S2) and expanding the ratio in $T$ (rather than separately expanding numerator and denominator) gives, to $O(T)$,

$$\beta_1 \approx -\frac{\alpha}{m + \alpha N(0)}\left[1 - \left(\frac{m}{2} + \alpha N(0)\right) T\right]. \tag{S40}$$

Higher baseline mortality $m$ weakens CNDD (less negative $\beta_1$) through two pathways: it dilutes the relative contribution of density-dependent mortality to total mortality risk at any instant, and it depletes the density field more rapidly over the interval. The $\alpha N(0)$ term inside the bracket reflects a third, related effect: density-dependent mortality itself thins the cohort, and this self-thinning further erodes the density signal as the interval lengthens. The transition rate $g_0$ does not enter $\beta_1$ at this order.

##### 5.3 Case 2: Density suppresses growth, density-independent mortality

**Assumptions.**

$$\mu_1 = m_1, \qquad \mu_2 = m_2, \qquad g(J_1, J_2) = g_0 \exp(-\gamma(J_1 + J_2)).$$

With density-independent mortality hazards, the mean hazard is

$$\bar{m}(t) = \frac{m_1 J_1(t) + m_2 J_2(t)}{N(t)} = m_1 + (m_2 - m_1)\frac{J_2(t)}{N(t)}. \tag{S41}$$

Thus $\bar{m}(t)$ depends on $N(0)$ only through the stage composition $J_2(t)/N(t)$, and only if $m_1 \neq m_2$.

**Short-time dynamics of $J_2$.** At $t = 0$, $J_1(0) = N(0)$ and $J_2(0) = 0$, so $\dot{J}_2(0) = g(0) \cdot N(0) = g_0\, e^{-\gamma N(0)}\, N(0)$. Therefore

$$J_2(t) = g_0\, e^{-\gamma N(0)}\, N(0)\, t + O(t^2).$$

Since $N(t) = N(0) + O(t)$,

$$\frac{J_2(t)}{N(t)} = g_0\, e^{-\gamma N(0)}\, t + O(t^2). \tag{S42}$$

**Cumulative hazard and sensitivity.**

$$\mathrm{H}(T) = m_1 T + (m_2 - m_1) \int_0^T \frac{J_2(t)}{N(t)}\, dt = m_1 T + \frac{1}{2}(m_2 - m_1)\, g_0\, e^{-\gamma N(0)}\, T^2 + O(T^3).$$

Differentiating with respect to $N(0)$,

$$\frac{\partial \mathrm{H}(T)}{\partial N(0)} = -\frac{1}{2}(m_2 - m_1)\, g_0\, \gamma\, e^{-\gamma N(0)}\, T^2 + O(T^3).$$

Expanding $e^{-\gamma N(0)} \approx 1$ at leading order in $\gamma N(0)$,

$$\frac{\partial \mathrm{H}(T)}{\partial N(0)} \approx \frac{1}{2}\, g_0\, \gamma\, (m_1 - m_2)\, T^2 + O(T^3). \tag{S43}$$

There is no $O(T)$ term: density acts through growth suppression rather than directly on mortality, so there is no instantaneous survival consequence at leading order. The sign depends on the mortality contrast: if $m_1 > m_2$ (mortality decreases with size, as is typical for seedlings), growth suppression retains individuals longer in the high-hazard small stage, strengthening realized interval CNDD.

**Regression coefficient.** At leading order, $\mathrm{H}(T) \approx m_1 T$. Substituting this and Eq. S43 into Eq. S2,

$$\beta_1 \approx -\frac{1}{2}\, g_0\, \gamma \left(1 - \frac{m_2}{m_1}\right) T. \tag{S44}$$

Higher baseline small-stage mortality $m_1$ strengthens CNDD (more negative $\beta_1$), since the mortality contrast $1 - m_2/m_1$ grows toward 1 as $m_1$ increases relative to $m_2$, preserving more of the size-dependent mortality difference that drives the effect. $\beta_1$ also grows in magnitude with census interval length $T$, reflecting the gradual build-up of density-dependent effects through growth suppression.

#### 5.4 Case 3: Large individuals drive density-dependent mortality of small individuals

**Assumptions.**

$$g(J_1, J_2) = g_0, \qquad \mu_1 = m_1 + \alpha J_2, \qquad \mu_2 = m_2.$$

Only small individuals are affected by crowding, and only large individuals generate it. Since $J_2(0) = 0$, there is no density effect at $t = 0$.

**Mean hazard.**

$$\bar{m}(t) = \frac{(m_1 + \alpha J_2)J_1 + m_2 J_2}{N} = \frac{m_1 J_1 + m_2 J_2}{N} + \alpha \frac{J_1 J_2}{N}.$$

The first term does not depend on $N(0)$, so

$$\frac{\partial \mathrm{H}(T)}{\partial N(0)} = \alpha \frac{\partial}{\partial N(0)} \int_0^T \frac{J_1(t) J_2(t)}{N(t)} \, dt. \tag{S45}$$

**Short-time evaluation.** Since the integrand already carries a factor $\alpha$, we evaluate it at leading order using the $O(t)$ expansions of $J_1$ and $J_2$:

$$J_1(t) = N(0)\,(1 - (m_1 + g_0)t) + O(t^2), \qquad J_2(t) = g_0 N(0)\, t + O(t^2), \qquad N(t) = N(0) + O(t).$$

Therefore

$$\frac{J_1 J_2}{N} = g_0 N(0)\, t + O(t^2), \qquad \int_0^T \frac{J_1 J_2}{N} \, dt = \frac{1}{2} g_0 N(0)\, T^2 + O(T^3).$$

**Cumulative hazard and sensitivity.**

$$\mathrm{H}(T) = \int_0^T \bar{m}(t)\, dt = \int_0^T \frac{m_1 J_1 + m_2 J_2}{N} \, dt + \frac{1}{2} \alpha g_0 N(0)\, T^2 + O(T^3).$$

Differentiating with respect to $N(0)$,

$$\frac{\partial \mathrm{H}(T)}{\partial N(0)} \approx \frac{1}{2} \alpha\, g_0\, T^2 + O(T^3). \tag{S46}$$

There is no $O(T)$ term because the density-generating stage ($J_2$) is absent at $t = 0$.

**Regression coefficient.** At leading order, $\mathrm{H}(T) \approx m_1 T$. Substituting this and Eq. S46 into Eq. S2,

$$\beta_1 \approx -\frac{\alpha\, g_0}{2m_1}\, T. \tag{S47}$$

Faster transition $g_0$ strengthens CNDD (more negative $\beta_1$) by accelerating accumulation of the large-individual density field, allowing the density-dependent mechanism to build up sooner. Higher baseline small-stage mortality $m_1$ weakens $\beta_1$ because small individuals die before large individuals have had time to accumulate and generate density-dependent pressure.

#### 5.5 Case 4: Small individuals generate and experience density-dependent mortality

**Assumptions.**

$$g(J_1, J_2) = g_0, \qquad \mu_1 = m_1 + \alpha J_1, \qquad \mu_2 = m_2.$$

The density effect is present immediately at $t = 0$ since $J_1(0) = N(0) > 0$.

**Mean hazard.**

$$\bar{m}(t) = \frac{(m_1 + \alpha J_1)J_1 + m_2 J_2}{N} = \frac{m_1 J_1 + m_2 J_2}{N} + \alpha \frac{J_1^2}{N},$$

so

$$\frac{\partial \mathrm{H}(T)}{\partial N(0)} = \alpha \, \frac{\partial}{\partial N(0)} \int_0^T \frac{J_1(t)^2}{N(t)}\, dt. \tag{S48}$$

**Short-time expansion of the integrand.** Writing $J_1^2/N = J_1 \cdot (J_1/(J_1 + J_2))$ and using

$$\frac{J_1}{J_1 + J_2} \approx 1 - g_0 t + O(t^2),$$

together with the short-time expansion of $J_1$ under the full (DD) dynamics,

$$\dot{J}_1(0) = -(g_0 + m_1 + \alpha N(0))N(0) \quad \Rightarrow \quad J_1(t) = N(0) - (g_0 + m_1 + \alpha N(0))N(0)\, t + O(t^2),$$

we obtain

$$\frac{J_1^2}{N} = N(0) - (m_1 + 2g_0 + \alpha N(0))N(0)\, t + O(t^2).$$

**Cumulative hazard and sensitivity.**

$$\mathrm{H}(T) = \int_0^T \bar{m}(t)\, dt = \int_0^T \frac{m_1 J_1 + m_2 J_2}{N}\, dt + \alpha \left[ N(0)\, T - \frac{1}{2}(m_1 + 2g_0 + \alpha N(0))N(0)\, T^2 \right] + O(T^3).$$

Differentiating with respect to $N(0)$,

$$\frac{\partial \mathrm{H}(T)}{\partial N(0)} = \alpha T - \left( \frac{\alpha m_1}{2} + \alpha g_0 + \alpha^2 N(0) \right) T^2 + O(T^3). \tag{S49}$$

Unlike Case 3, there is an $O(T)$ term because the density-generating stage is present at $t = 0$.

**Regression coefficient.** At leading order, $\mathrm{H}(T) \approx (m_1 + \alpha N(0))T$. Substituting this and Eq. S49 into Eq. S2 and expanding the ratio in $T$ gives, to $O(T)$,

$$\beta_1 \approx -\frac{\alpha}{m_1 + \alpha N(0)} \left[ 1 - \left( \frac{m_1}{2} + g_0 + \alpha N(0) \right) T \right]. \tag{S50}$$

Consistent with Case 1, larger $m_1$ weakens realized interval CNDD, both by diluting the relative importance of density-dependent mortality and through the self-thinning captured by the $\alpha N(0)$ term. Unlike Cases 2 and 3, increasing $g_0$ also weakens realized interval CNDD by accelerating escape from the small stage where density dependence is strongest.

### 6 Instantaneous density sensitivity and realized interval CNDD

The main text notes that, unsurprisingly, greater instantaneous density sensitivity ($\alpha$ or $\gamma$) steepens the relationship between density-independent vital rates and realized interval CNDD (see main text, "How do density-independent vital rates shape CNDD?"). Here we confirm this directly, using the same closed-cohort size-structured model underlying Fig. 3 and Box 1, holding baseline vital rates fixed at representative Case 1 and Case 2 values and sweeping the focal sensitivity parameter.

For density-dependent mortality ($\alpha > 0$), we use Case 1's symmetric mortality setup, with baseline mortality $m_0$ fixed at a modest value and $\alpha$ swept across several orders of magnitude (Fig. S2A,C,E). For density-dependent growth suppression, we use Case 2's setup, with $m_0$ fixed above $m_{\min}$ (as required for Case 2's mortality-contrast mechanism to operate at all) and $\gamma$ swept analogously (Fig. S2B,D,F). In both cases, increasing the focal sensitivity parameter steepens the decline of survival with initial density (Fig. S2A–B) and makes $\beta_1$ progressively more negative (Fig. S2C–D), confirming that instantaneous density sensitivity strengthens realized interval CNDD as expected, independently of the baseline-rate effects that are the main text's focus. The corresponding density-dependent hazard increment (Fig. S2E–F) shows this directly: higher $\alpha$ or $\gamma$ produces a larger and more persistent excess hazard over the census interval, particularly at high initial density, mirroring the panel-I–L logic of Fig. 3 but here driven by instantaneous sensitivity rather than baseline demography.

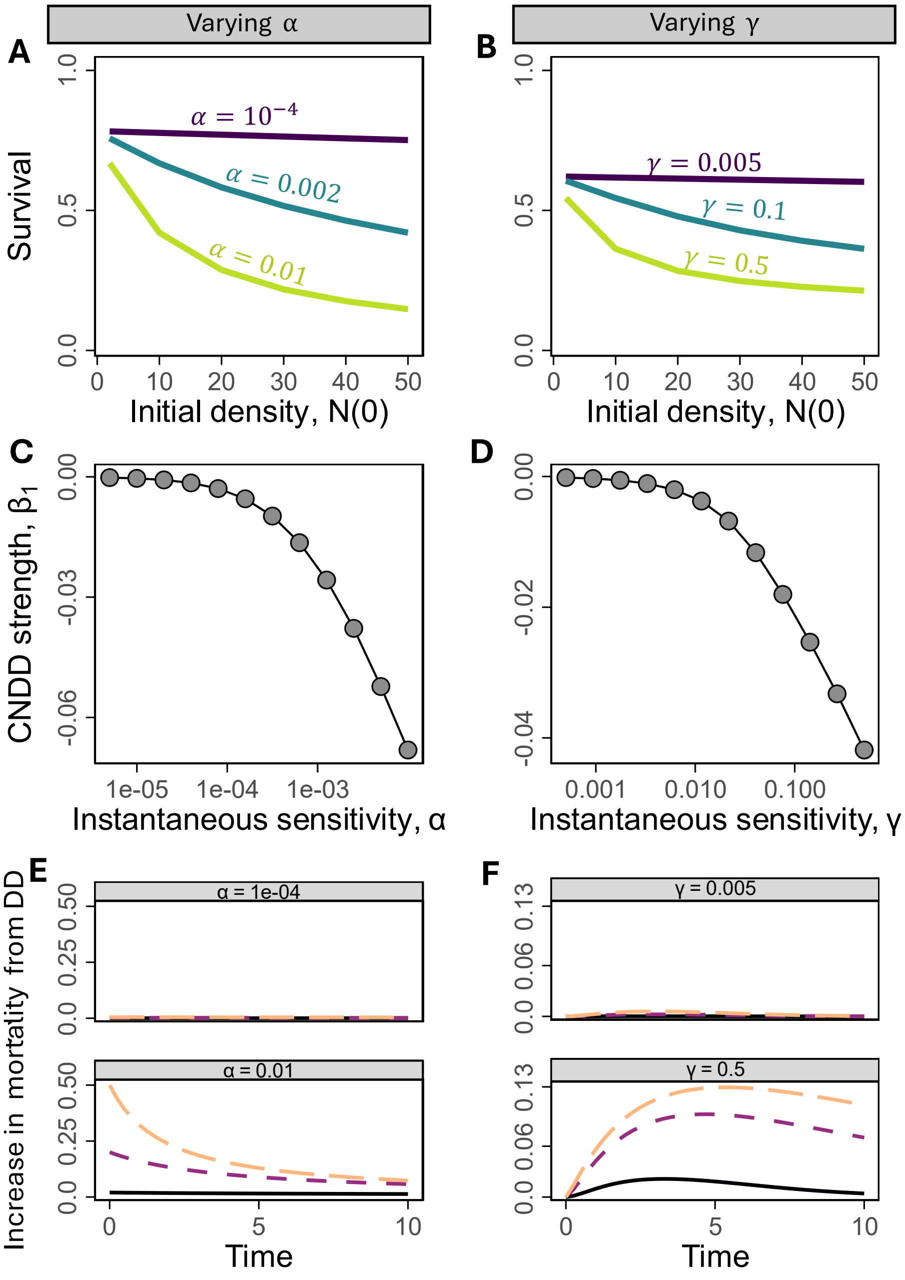

Figure S2: **Greater instantaneous density sensitivity steepens realized interval CNDD.** Left column: mortality channel (Case 1 background), sweeping $\alpha$. Right column: growth-suppression channel (Case 2 background), sweeping $\gamma$. (A–B) Survival probability $\exp(-\mathrm{H}(T))$ at interval time $T$ as a function of initial cohort size $N(0)$, for three values of the focal sensitivity parameter. (C–D) Measured CNDD strength, summarized as $\beta_1$ — the logit-scale density slope (Eq. 14 from the main text) — as a function of the focal sensitivity parameter, on a log scale. More negative values indicate stronger CNDD. (E–F) The density-dependent increment to the instantaneous per-capita hazard over the census interval, shown for a low (bottom facet) and high (top facet) value of the focal parameter, and for three initial cohort sizes (line colors; see legend in Fig. 3). All other parameters held fixed within each column; see Table X for all parameters.

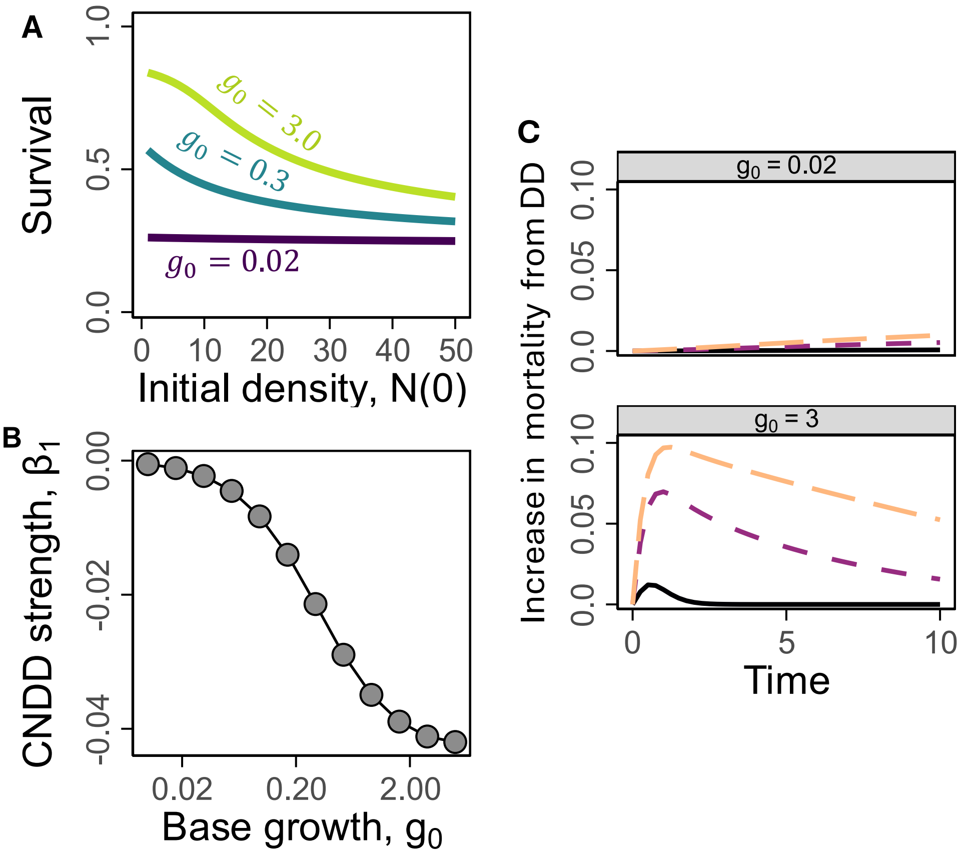

Figure S3: **Faster baseline growth strengthens CNDD in Case 2.** Baseline mortality is fixed at $m_0 = 0.15$ throughout (above $m_{\min} = 0.01$, so the Case 2 mortality-contrast mechanism is active). (A) Survival probability $\exp(-\mathrm{H}(T))$ as a function of initial cohort size $N(0)$, for $g_0 = 0.02$, 0.3, and 3.0 (colors). (B) Measured CNDD strength ($\beta_1$) as a function of $g_0$, swept from 0.01 to 5 (log scale). (C) The density-dependent increment to the instantaneous per-capita hazard over the census interval, shown for $g_0 = 0.02$ (top facet) and $g_0 = 3.0$ (bottom facet), for initial cohort sizes $N(0) = 2$, 20, and 50 (line colors).

### 7 Case 2: the effect of baseline growth $g_0$

In Case 2, density suppresses the transition rate from the small to the large size class ($\gamma > 0$), with mortality varying by size ($m_1 > m_2$; main text; Box 1). Box 1 shows analytically that, in addition to strengthening with the mortality contrast between stages, CNDD also strengthens with faster baseline growth $g_0$ (Eq. B6). Here we confirm this numerically, using the same closed-cohort size-structured model underlying Fig. 3 and Box 1, with parameters otherwise matching Fig. 3B,F,J (Table S1) except $g_0$ swept as the focal parameter (Fig. S3).

### 8 Equilibrium analysis of PDE model

Throughout the main text, we analyze how density-independent vital rates shape realized interval CNDD within closed cohorts, where the relevant baseline rates act by altering the dynamics of $P(t)$ and cohort composition over a finite interval $[0, T]$. The same demographic mechanisms operate at equilibrium ($f > 0$), but through a different pathway: rather than shaping how $P(t)$ and the size distribution evolve within an interval, baseline rates determine the standing crowding field $P^*$ and equilibrium size distribution $n^*(x)$ themselves. Here we show numerically that the same four demographic scenarios introduced in Box 1 yield qualitatively identical effects on realized interval CNDD when evaluated at equilibrium.

#### 8.1 Model and equilibrium definition

We use the equilibrium variant of the size-structured PDE model introduced in the main text (Methods: PDE framework), in which seed input $f > 0$ sustains a standing seedling population rather than a single closed cohort. Equilibrium corresponds to a steady state of the model, $\partial n(x,t)/\partial t = 0$, in which the seed pool, size distribution, and crowding field $P(t)$ no longer change in time. Because the crowding-dependent terms in Eqs. 5–6 in the main text enter through $P(t)$, which is itself an integral over the size distribution $n(x,t)$ (Eq. 7, main text), the equilibrium size distribution $n^*(x)$ and the equilibrium crowding field $P^*$ must be solved for jointly: $n^*(x)$ depends on $P^*$, and $P^*$ is in turn defined as an integral of $n^*(x)$. This does not admit a general closed-form solution, so we characterize $n^*(x)$, $P^*$, and their dependence on baseline vital rates numerically, by integrating the model forward in time until it converges to a steady state.

#### 8.2 "Tagged" cohort approach

To connect the equilibrium size distribution to realized interval CNDD, we adapt the hazard-based framework used throughout the main text (Methods: Hazard-based metrics) to the equilibrium setting. At a genuine numerical equilibrium for given values of the baseline vital rate parameters, we conceptually divide the standing population into two groups: individuals present at the moment of a hypothetical census, and individuals recruited afterward. Both groups continue to grow and experience mortality according to Eqs. 5–6, and both contribute to a single, shared crowding field $P(t)$, since real conspecific crowding does not distinguish between previously established and newly arrived individuals. Only the originally censused group, however, contributes to the cumulative hazard $\mathrm{H}(T)$ and survival $\exp(-\mathrm{H}(T))$ that we report, exactly as an empirical census study would track survival of a "tagged" group of individuals over a subsequent interval while new recruits continue to arrive and contribute to local density.

This approach allows us to compute $\mathrm{H}(T)$, and hence $\beta_1$ via Eq. S2, starting from a population that is already at equilibrium rather than from a freshly established cohort, while still allowing recruitment to continue exactly as it does at equilibrium. We repeat this procedure across a range of equilibrium densities for each focal parameter value, obtaining survival as a function of initial density and the corresponding $\beta_1$, directly analogous to the closed-cohort results presented in Fig. 3 and Box 1.

#### 8.3 Results

Across all four demographic scenarios introduced in Box 1, the equilibrium analysis recovers the same qualitative relationship between baseline vital rates and realized interval CNDD found in the closed-cohort analysis (Fig. S4). Lower baseline mortality $m_0$ strengthens $\beta_1$ when density acts through mortality directly (Case 1; Fig. S4A,E), while higher $m_0$ strengthens $\beta_1$ when density acts through growth suppression (Case 2; Fig. S4B,F), mirroring the dependence on baseline mortality identified in the closed-cohort cases. Likewise, faster baseline growth $g_0$ strengthens $\beta_1$ when density-dependent effects are concentrated in large individuals (Case 3; Fig. S4C,G), and weakens $\beta_1$ when those effects are concentrated in small individuals (Case 4; Fig. S4D,H), again matching the corresponding closed-cohort result. These results confirm that the demographic mechanisms described in the main text are not specific to closed cohorts, but extend to standing, equilibrium populations maintained by ongoing recruitment.

The demographic mechanisms underlying these results differ in their relationship to the closed-cohort cases.

In Case 1, the closed-cohort mechanism combines dilution of the relative contribution of density-dependent mortality with depletion of the cohort over the census interval. At equilibrium, an analogous depletion still operates: a cohort tracked forward from a standing equilibrium thins out over the tracked interval exactly as a closed cohort does, and does so more slowly when baseline mortality $m_0$ is lower, so more of the cohort survives to keep accumulating crowding-driven hazard. This is the same fact as the dilution argument viewed differently: at any instant, lower $m_0$ means density-dependent mortality constitutes a larger share of total hazard, and this is what makes the larger surviving fraction translate into a larger accumulated excess, $\mathrm{H}_{\mathrm{DD}}(T)$, reproducing the same direction of effect on $\beta_1$.

In Case 2, the mechanism transfers directly: because density suppresses growth rather than acting on mortality, realized interval CNDD depends on the mortality contrast between small and large size classes $(1 - m_{\min}/m_0)$, and this contrast is a property of the baseline rates themselves, not something that builds up over an interval, so it operates identically at equilibrium.

In Case 3, the closed-cohort mechanism depends on how quickly large, crowding-generating individuals accumulate from an initial cohort with none present; at equilibrium there is no such accumulation, and the relevant quantity instead becomes the standing abundance of large individuals already present in the equilibrium size distribution $n^*(x)$, which itself increases with $g_0$. Faster baseline growth therefore strengthens $\beta_1$ at equilibrium by shifting the standing size distribution toward larger individuals, rather than by accelerating their accumulation within the census interval.

Case 4 reflects the closed-cohort escape-rate mechanism directly: faster growth $g_0$ continues to move individuals out of the small, crowding-affected stage during the census interval itself, shortening their exposure to density-dependent mortality and attenuating realized interval CNDD, the same mechanism identified in the closed-cohort case.

Because the equilibrium size distribution and crowding field have no general closed-form solution, the results above are characterized numerically rather than derived analytically, in contrast to the closed-cohort results in Box 1. For Cases 3 and 4, in which baseline growth $g_0$ is the focal parameter, we restrict the range of $g_0$ considered to values for which the equilibrium size distribution remains numerically well-behaved; at sufficiently high $g_0$, individuals can reach the upper boundary of the size domain faster than the discretization used here resolves, producing a numerical artifact unrelated to the underlying biology.

#### 8.4 Equilibrium size distributions

To illustrate how baseline growth reshapes the standing population at equilibrium, we compare the equilibrium size distribution $n^*(x)$ for two values of baseline growth $g_0$ in Case 3 and Case 4 (Fig. S5), the two cases in which $g_0$ is the focal parameter. For each case, we bisect the recruitment rate $f$ separately for each $g_0$ value so that both reach the same equilibrium density $N^*$, isolating the effect of $g_0$ on the shape of $n^*(x)$ from any accompanying change in standing density. In both cases, faster baseline growth shifts $n^*(x)$ toward larger individuals, consistent with the mechanism described above: a larger share of the standing population occupies large size classes when growth is faster, regardless of whether crowding is generated by large individuals (Case 3) or by small individuals (Case 4). This shift in $n^*(x)$ is the basis for the equilibrium results reported for Cases 3 and 4 in the main text and above.

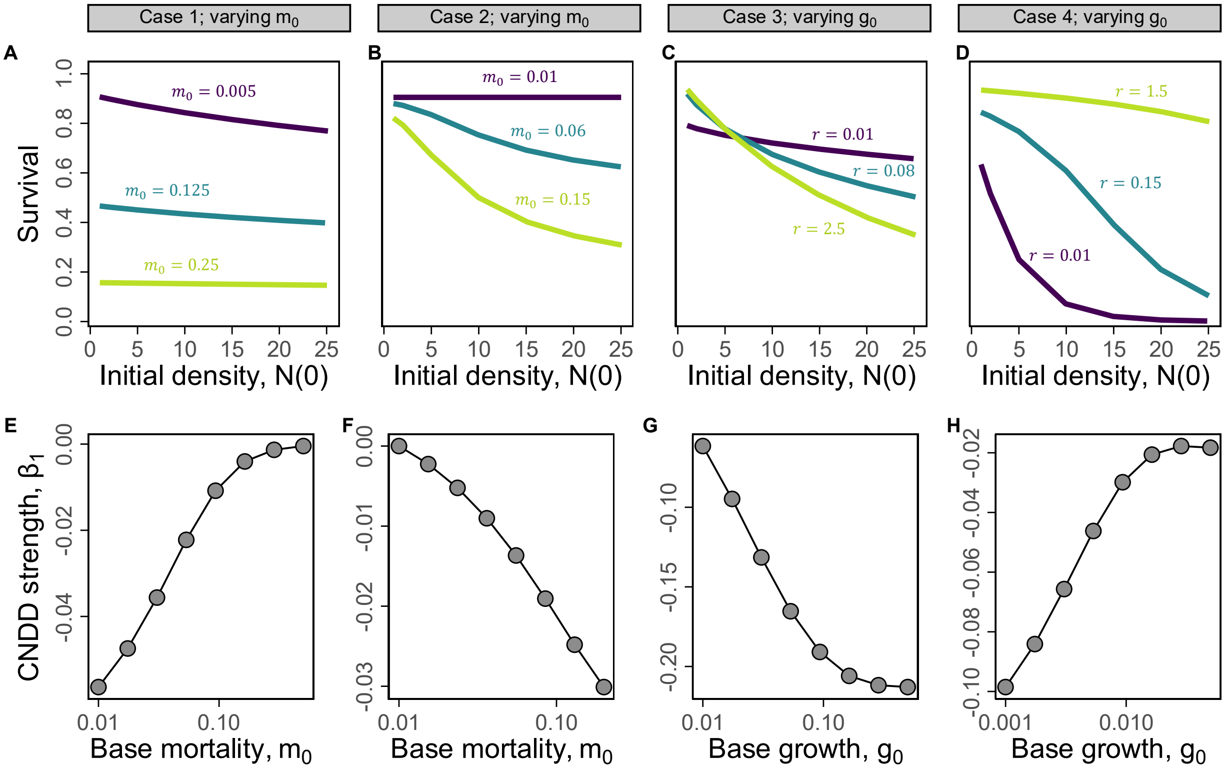

Figure S4: **Density-independent vital rates modulate CNDD at equilibrium.** Each column shows the same demographic scenario as the corresponding case in Fig. 3 and Box 1 (Cases 1–4), but evaluated at equilibrium ($f > 0$) rather than within a closed cohort. (A–D) Survival of a cohort tagged at a numerical equilibrium and tracked forward over a census interval $T$, as a function of the equilibrium density $N^*$ at which it was tagged, for three values of the focal density-independent parameter (colors; darker shades indicate lower parameter values). For each parameter value, the recruitment rate $f$ was chosen separately for each plotted $N^*$ so that the underlying population was at its own genuine equilibrium, rather than an initial condition perturbed away from one; recruitment continued throughout the tracked interval, with new recruits sharing the same crowding field as the tagged cohort but excluded from the survival outcome shown. (E–H) Measured CNDD strength, summarized as $\beta_1$ — the exact logit-scale density slope (Eq. 14, main text) — as a function of the focal parameter; more negative values indicate stronger CNDD. $\beta_1$ is computed using the same central-difference approach as Fig. 3E–H, evaluated at each of several equilibrium densities $N^*$ and averaged across them. Columns correspond to: (A,E) varying baseline small-size mortality $m_0$ when density acts through mortality ($\alpha > 0$, $\gamma = 0$); (B,F) varying $m_0$ when density acts through growth suppression ($\gamma > 0$, $\alpha = 0$); (C,G) varying baseline growth rate $g_0$ when density-dependent effects increase with individual size; and (D,H) varying $g_0$ when density-dependent effects are concentrated in small individuals. All other parameters are held fixed within each column and match the corresponding column in Fig. 3 (see Table S2 for full parameter values).

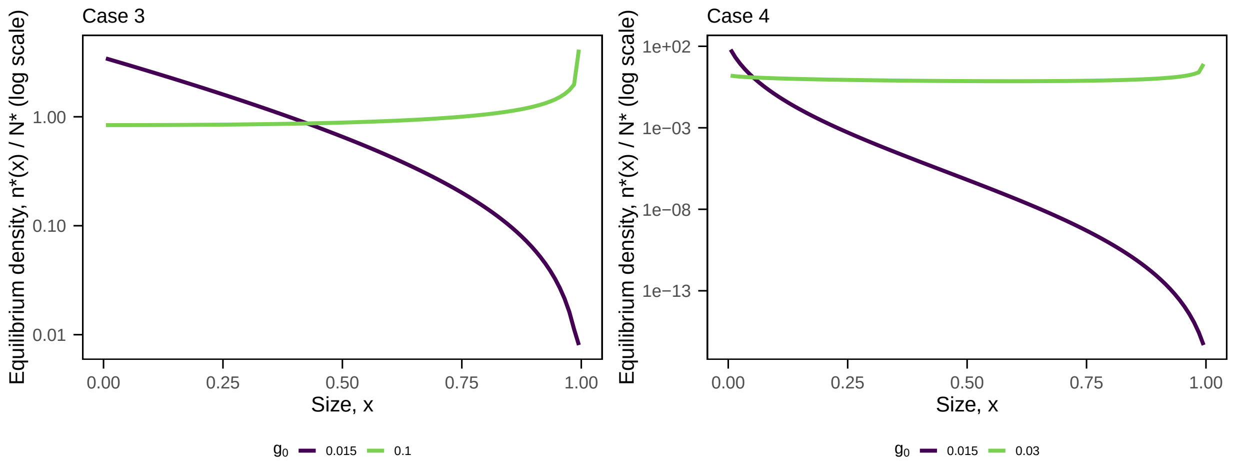

Figure S5: **Faster baseline growth shifts the equilibrium size distribution toward larger individuals.** Equilibrium size distribution $n^*(x)$ (normalized so that $\int n^*(x)\,dx = 1$; log scale) for two values of baseline growth $g_0$ in Case 3 (left; large individuals generate density-dependent pressure, $p$-weight$= 1.0$) and Case 4 (right; small individuals generate and experience density-dependent pressure, $p$-weight$= -0.8$), the two cases in which $g_0$ is the focal parameter. For each $g_0$ value, the recruitment rate $f$ was chosen so that the equilibrium density matched $N^* = 25$ across both curves within a panel, isolating the effect of $g_0$ on the shape of $n^*(x)$ from any accompanying change in standing density. In both cases, faster baseline growth ($g_0 = 0.1$ in Case 3; $g_0 = 0.03$ in Case 4) shifts mass away from the smallest size classes and toward larger individuals, relative to the slower-growth curve ($g_0 = 0.015$ in both panels).

### 9 Finite-volume approximation and mean-field limit of the Gillespie SSA

The spatially explicit multispecies simulations (Methods: Spatially explicit multispecies simulations) track integer abundances $N_{s,\ell}(t)$ across discrete size bins using a Gillespie SSA. Here we show that the mean-field limit of this event-based formulation recovers a finite-volume ODE approximation of the size-structured PDE (Eq. 4 of the main text).

Size is discretized on $[0, x_{\max}]$ into $b$ equal bins of width $\Delta x = x_{\max}/b$, with bin $\ell$ spanning $[(\ell-1)\Delta x,\ \ell\Delta x]$ and midpoint $x_\ell = (\ell - \frac{1}{2})\Delta x$ ($\ell = 1, \ldots, b$); individuals in bin $\ell$ are treated as located at $x_\ell$. Growth events move one individual from bin $\ell$ to bin $\ell + 1$ at per-capita rate $g(x_\ell, t)/\Delta x$, and mortality events remove one individual from bin $\ell$ at per-capita rate $m(x_\ell, t)$. Taking expectations over the stochastic process, the mean-field dynamics of $N_{s,\ell}(t)$ are

$$\frac{dN_{s,\ell}}{dt} = \frac{g(x_{\ell-1}, t)}{\Delta x} N_{s,\ell-1} - \frac{g(x_\ell, t)}{\Delta x} N_{s,\ell} - m(x_\ell, t)\, N_{s,\ell}, \quad \ell = 1, \ldots, b, \tag{S51}$$

with $N_{s,0} \equiv 0$ (no inflow at the lower boundary in closed-cohort analyses). Setting $n_\ell(t) = N_{s,\ell}(t)/\Delta x$ as the bin density, this becomes

$$\frac{dn_\ell}{dt} = \frac{g(x_{\ell-1}, t)\, n_{\ell-1} - g(x_\ell, t)\, n_\ell}{\Delta x} - m(x_\ell, t)\, n_\ell, \tag{S52}$$

which is the standard upwind finite-volume discretization of the advection–mortality PDE from the main text,

$$\frac{\partial n}{\partial t} + \frac{\partial}{\partial x}\left[g(x, t)\, n\right] = -m(x, t)\, n, \tag{S53}$$

on the grid $\{x_\ell\}$. As $\Delta x \to 0$ (equivalently $b \to \infty$), the finite-volume scheme converges to the PDE under standard smoothness conditions on $g$ and $m$ (e.g., LeVeque, 2002). The Gillespie SSA therefore provides an exact stochastic realization of the individual-level process whose mean-field limit is the size-structured PDE.

### 10 Spatially explicit multispecies simulations and GLMM fitting details

We provide here a more detailed description of how census-interval survival datasets are constructed from Gillespie SSA simulations, and of the GLMM fitting procedure applied to those datasets.

#### 10.1 Simulation landscape and community initialization

Simulations are run on a toroidal $n \times n$ grid of discrete patches. Each patch $j$ is assigned a habitat value $E_j$ drawn from a spatially autocorrelated Gaussian field: an i.i.d. $N(0, 1)$ field is convolved with a Gaussian smoothing kernel on the torus, then standardized to mean 0 and unit variance. The smoothing bandwidth is set to produce a target spatial autocorrelation (parameter $= 0.6$ in all simulations reported here).

Adults are placed on the grid at initialization and held fixed throughout the simulation, representing a static overstory that provides a continuous seed source. A total of $\lfloor \text{occupancy} \times n^2 \rfloor$ adult slots are allocated across all species (occupancy $= 0.25$ in all simulations). Species abundances are drawn from a lognormal distribution with log-scale standard deviation $\sigma_{\log} = 1.2$, subject to a minimum count of 10 adults per species; the resulting counts are then normalized to sum to the total adult slots. Spatial positions of each species' adults are assigned by placing individuals sequentially into unoccupied patches, biased toward patches with high values of a species-specific spatially autocorrelated random field (using the same smoothing bandwidth as the habitat field), producing realistic within-species spatial aggregation.

Seed dispersal from adults to patches follows a Gaussian kernel truncated at radius $r_{\text{kern}}$: the seed rain into patch $j$ from species $s$ is proportional to $\sum_{j'} A_{s,j'} \exp(-d_{jj'}^2/2\sigma_{\text{kern}}^2)$, where $A_{s,j'}$ is the number of adults of species $s$ in patch $j'$, $d_{jj'}$ is the toroidal distance between patches, $\sigma_{\text{kern}} = 3$ patches, and $r_{\text{kern}} = 12$ patches. The seed rain field is precomputed at initialization and held fixed. Within each patch, seeds arrive at a Poisson rate proportional to the precomputed seed rain scaled by species fecundity, are lost at per-capita rate $m_B$, and germinate at per-capita rate $g_B$, entering the smallest size bin as new seedlings.

#### 10.2 Census interval generation

The Gillespie SSA evolves a continuous-time Markov chain over discrete size bins, tracking integer seedling counts $N_{s,\ell}(t)$ and an integer seed pool $S_s(t)$ for each species $s$ and size bin $\ell$ within each patch. Rather than recording individual trajectories directly—which would be prohibitively expensive—the simulation maintains a list of unique integer identifiers (`local_id`) for each seedling currently occupying each bin. Identifiers are assigned sequentially at germination: when a germination event occurs for species $s$, a new seedling enters bin 1 and receives the next available integer ID. When a growth event occurs in bin $(s, \ell)$, one of the $N_{s,\ell}(t)$ currently resident IDs is selected uniformly at random and transferred to bin $\ell + 1$. When a death event occurs in bin $(s, \ell)$, one ID is selected uniformly at random and permanently removed, with the event recorded in a death log. This random-selection scheme is equivalent to assuming all individuals within a species–bin cell are exchangeable, consistent with the mean-field model.

Census snapshots are recorded at fixed intervals of length $T$ (the census interval). At each snapshot, the full list of IDs present in each species–bin cell is written to the output, along with the patch environment $E_j$. From these records, the number of conspecific and heterospecific individuals present in each patch at each census time is computed by counting IDs across all size bins, and used as the raw crowding covariates in the GLMM (see below).

From consecutive census snapshots separated by $T$, we construct individual-level survival records in R. For each individual $i$ present at census time $t_c$, we define the binary outcome $y_i = 1$ if the same `local_id` appears in the subsequent census at $t_c + T$, and $y_i = 0$ otherwise (i.e., if the ID is absent, indicating death during the interval). Covariates are measured at the *start* of the interval (census time $t_c$): initial size $\text{Size}_i$ is the midpoint of the size bin occupied at $t_c$; conspecific crowding $\mathrm{Con}_{i,\mathrm{raw}} = P_{\mathrm{con},s(i)}(t_c)$ and heterospecific crowding $\mathrm{Het}_{i,\mathrm{raw}} = P_{\mathrm{het},s(i)}(t_c)$ are the crowding values recorded at $t_c$. To reduce transient effects from simulation initialization, only census intervals from the final $K$ census times are retained for analysis ($K = 20$ in Figs. 4 and 5).

#### 10.3 GLMM fitting procedure

Raw crowding covariates are the counts of conspecific and heterospecific seedling neighbors in the patch at the start of the census interval (excluding the focal individual), denoted $\mathrm{Con}_{i,\mathrm{raw}}$ and $\mathrm{Het}_{i,\mathrm{raw}}$ respectively. These are power-transformed and z-scored prior to fitting: $\mathrm{Con}_i = z\big(\mathrm{Con}_{i,\mathrm{raw}}^{b_1}\big)$ and $\mathrm{Het}_i = z\big(\mathrm{Het}_{i,\mathrm{raw}}^{b_2}\big)$, where $z(\cdot)$ denotes standardization to mean zero and unit variance within the fitted dataset and $(b_1, b_2)$ are power-transform exponents. Initial size $\mathrm{Size}_i$ and, where applicable, patch environment $E_{j(i)}$ are similarly z-scored.

The exponents $(b_1, b_2)$ are selected independently for each scenario by a grid search over $\{0.25, 0.5, 0.75, 1, 1.25, 1.5, 2\}^2$ (49 candidate pairs). For computational tractability, the grid search uses a random subsample of 600 patches and the Laplace approximation (`nAGQ = 0` in lme4). The pair maximizing the fitted log-likelihood is selected, and the model is then refit on the full dataset using adaptive Gauss–Hermite quadrature with one quadrature point (`nAGQ = 1`). For the variance decomposition (Fig. 5I), the exponents are selected from the "both vary" condition only and applied identically to all four conditions, ensuring that differences in $\widehat{\beta}_{\mathrm{con},s}$ across conditions reflect differences in the underlying data rather than differences in the crowding scale. The GLMM formula follows Eq. 9 in the main text, with correlated random intercepts and random slopes for both conspecific and heterospecific density by species: `(1 + Con + Het | species)`. Models were fit using lme4 (Bates et al., 2015) with the `bobyqa` optimizer (maximum $5 \times 10^4$ function evaluations).

Several fitted models produced singular random-effects covariance matrices (as flagged by `isSingular()` in lme4), arising consistently from near-zero estimated variance in the heterospecific density random slopes ($u_{\mathrm{het},s}$). This is expected: by simulation design, species differ primarily in their density-independent vital rates rather than in their sensitivity to heterospecific neighbors, so little genuine between-species variance in the heterospecific slope is available to estimate. Singular fits were accepted; because the degeneracy is confined to the heterospecific slope variance, point estimates of the conspecific density random slopes $u_{\text{con},s}$—the quantities of primary interest—remain well-identified. DHARMa residual diagnostics (Hartig, 2016) were performed on all final models; residuals were largely well-behaved, with minor deviations from uniformity in some scenarios.

### 11 GAM robustness analysis

Recent work on CNDD estimation has highlighted that the logit-linear relationship between conspecific density and survival assumed by GLMMs may be overly restrictive, and that flexible nonparametric approaches better capture the saturating nonlinearities common in empirical survival data (Hülsmann et al., 2024; Huanca-Nunez et al., 2026). Following Hülsmann et al. (2024), we complement our GLMM analysis with species-specific generalized additive models (GAMs) fit to the same simulated census datasets, confirming that the interspecific patterns in measured CNDD are robust to the choice of statistical model.

For each species $s$ in each scenario, we fit a separate GAM of the form

$$\log(-\log(1-\Pr(Y_{ij}=1))) = \beta_{0,s}+f_{\text{con},s}(\text{Con}_{ij,\text{raw}})+f_{\text{het},s}(\text{Het}_{ij,\text{raw}})+f_{\text{size},s}(\text{Size}_{ij})+u_j \quad \text{(S54)}$$

where $\Pr(Y_{ij}=1)$ is the predicted survival probability of individual $i$ in census interval $j$, the left-hand side is the complementary log-log (cloglog) link, $f_{\text{con},s}$, $f_{\text{het},s}$, and $f_{\text{size},s}$ are thin-plate regression splines with basis dimension $k=10$ (reduced when the number of unique values is less than $k$), and $u_j \sim N(0,\sigma_u^2)$ is a random intercept for census interval. The cloglog link was chosen for consistency with Hülsmann et al. (2024) and related works. Raw (untransformed) crowding counts were used as smooth predictors, with the GAM's flexible smooth handling nonlinearity directly. Models were fit by restricted maximum likelihood (REML) using the mgcv package (Wood and Wood, 2015) in R.

Species-level CNDD was quantified as the average marginal effect (aME) of conspecific density on survival probability (Hülsmann et al., 2024). For each observation $i$ of species $s$, we computed the predicted change in survival probability from adding one additional conspecific neighbor,

$$\text{aME}_{i,s} = p_{i,s}(\text{Con}_{i,\text{raw}}+1) - p_{i,s}(\text{Con}_{i,\text{raw}}),$$

where $p_{i,s} = \Pr(Y_{ij} = 1)$ is the predicted survival probability at the response scale, and averaged these across all observations for that species, $\mathrm{aME}_s = n_s^{-1} \sum_i \mathrm{aME}_{i,s}$. A more negative $\mathrm{aME}_s$ indicates stronger measured CNDD for species $s$. This quantity is analogous to the species-level random slope $\widehat{\beta}_{\mathrm{con},s}$ from the GLMM, but estimated nonparametrically and at the response (probability) rather than logit scale.

The resulting species-level aME estimates show the same qualitative relationship with the focal vital-rate parameter as the GLMM random slopes across all four simulation scenarios (Fig. S6): species with vital rates predicted by Box 1 to exhibit stronger measured CNDD tend to have more negative aME estimates, and vice versa. These results confirm that the interspecific patterns in measured CNDD reported in Fig. 5 are robust to the choice of statistical model and link function.

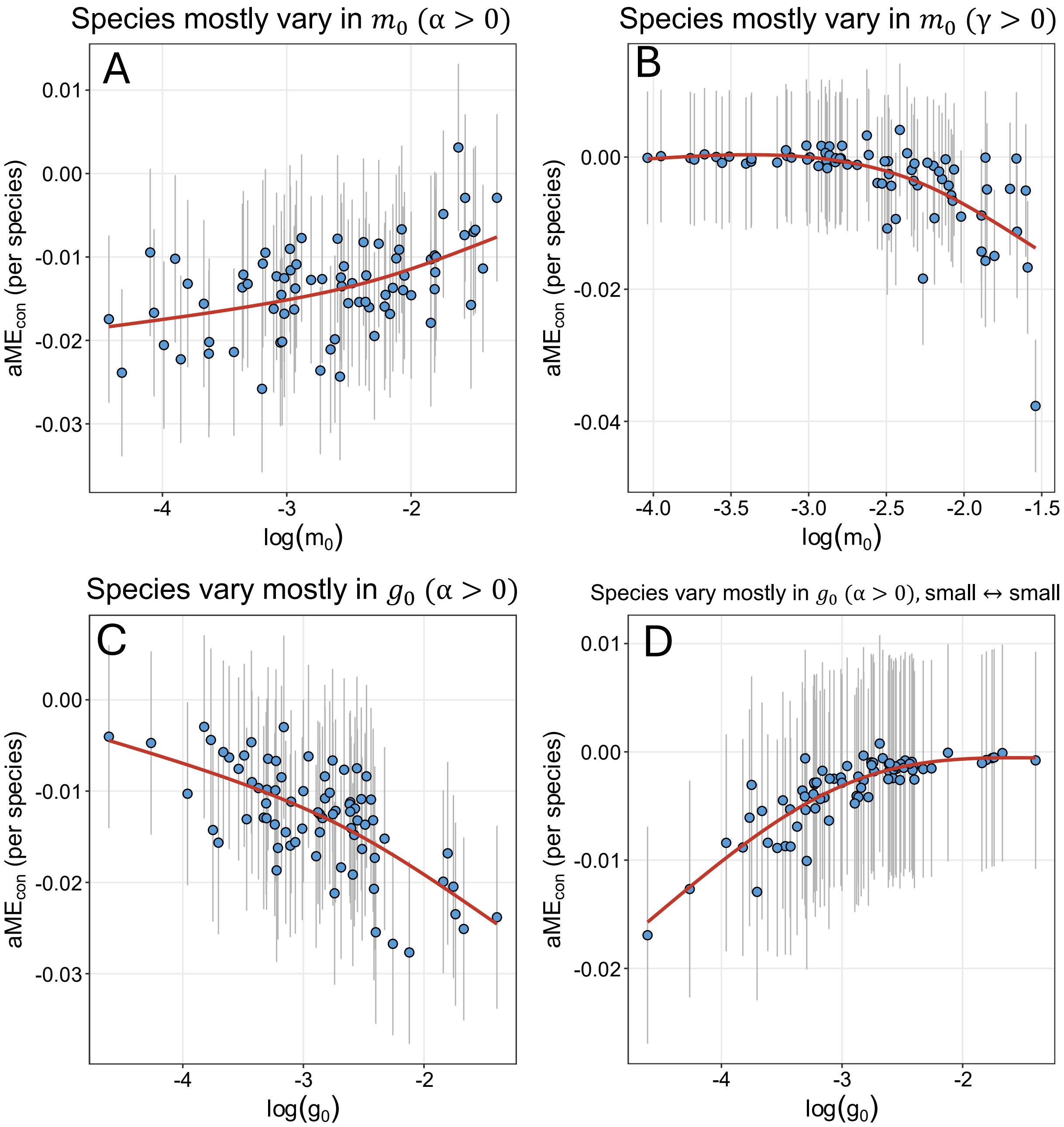

Figure S6: **GAM-based species-level CNDD estimates reproduce the patterns in Fig. 5.** Each panel corresponds to one demographic scenario from Fig. 5, refitted using species-specific generalized additive models (GAMs) with a complementary log-log link. The $y$-axis shows the average marginal effect (aME) of conspecific density on survival probability for each species: the mean predicted change in survival from adding one additional conspecific neighbour, averaged across all observations for that species. The $x$-axis shows the log of the focal vital-rate parameter. Error bars show the standard deviation of individual observation-level marginal effects within each species. The red line shows a GAM smooth of the relationship. In all four scenarios, the direction and pattern of the relationship between vital rates and species-level measured CNDD is qualitatively consistent with Fig. 5, confirming that results are robust to the choice of statistical model and link function.

### 12 Simulation parameters for Fig. 3, Fig. S2, and Fig. S4

We present parameters used in Fig. 3 in the main text, and Figs. S2 and S4 in the supplement.

Table S1: Parameters used in Figures 3, S2, and S3.

| Parameter | Value(s) |
|---|---|
| *Fig. 3A, E, I (Case 1: mortality channel, symmetric)* | |
| $T$ | 10 |
| $N(0)$ values | 2, 10, 20, 30, 40, 50 |
| $f$ | 0 |
| $g_0$ (baseline growth) | 0.25 |
| $m_{\min}$ | 0.01 |
| $\alpha$ | 0.0025 |
| $\gamma$ | 0 |
| $p$ | 0.0 |
| $q$ | 0.0 |
| $m_0$ | 0.005–3 |
| *Fig. 3B, F, J (Case 2: growth-suppression channel)* | |
| $T$ | 10 |
| $N(0)$ values | 2, 10, 20, 30, 40, 50 |
| $f$ | 0 |
| $g_0$ (baseline growth) | 0.5 |
| $m_{\min}$ | 0.01 |
| $\alpha$ | 0 |
| $\gamma$ | 0.3 |
| $p$ | 1.0 |
| $q$ | 0.0 |
| $m_0$ | 0.01–1.1 |
| *Fig. 3C, G, K (Case 3: large individuals drive DD mortality)* | |
| $T$ | 10 |
| $N(0)$ values | 2, 10, 20, 30, 40, 50 |
| $f$ | 0 |
| $m_{\min}$ | 0.001 |
| $m_0$ | 0.03 |
| $\alpha$ | 0.005 |
| $\gamma$ | 0 |
| $p$ | 1.0 (large neighbors dominate crowding) |
| $q$ | 0.0 |
| $g_0$ | 0.01–5 |
| *Fig. 3D, H, L (Case 4: small individuals generate and experience DD mortality)* | |
| $T$ | 10 |
| $N(0)$ values | 2, 10, 20, 30, 40, 50 |
| $f$ | 0 |
| $m_{\min}$ | 0.003 |
| $m_0$ | 0.03 |
| $\alpha$ | 0.0001 |
| $\gamma$ | 0 |
| $p$ | −0.8 (small neighbors dominate crowding) |
| $q$ | −0.8 (small individuals more susceptible) |

*(continued)*

| Parameter | Value(s) |
| --- | --- |
| $g_0$ | 0.02–5 |
| *Fig. S4A, C, E (mortality channel, Case 1 background, varying $\alpha$)* | |
| $T$ | 10 |
| $N(0)$ values | 2, 10, 20, 30, 40, 50 |
| $f$ | 0 |
| $g_0$ (baseline growth) | 0.25 |
| $m_0$ | 0.05 |
| $m_{\min}$ | 0.01 |
| $\gamma$ | 0 |
| $p$ | 0.0 |
| $q$ | 0.0 |
| $\alpha$ | 0.000005–0.01 |
| *Fig. S4B, D, F (growth-suppression channel, Case 2 background, varying $\gamma$)* | |
| $T$ | 10 |
| $N(0)$ values | 2, 10, 20, 30, 40, 50 |
| $f$ | 0 |
| $g_0$ (baseline growth) | 0.5 |
| $m_0$ | 0.2 |
| $m_{\min}$ | 0.01 |
| $\alpha$ | 0 |
| $p$ | 1.0 |
| $q$ | 0.0 |
| $\gamma$ | 0.0005–0.5 |
| *Fig. S4A, E (Case 1: mortality channel, varying $m_0$)* | |
| $N^*$ values | 1, 2, 5, 10, 15, 20, 25 |
| $f$ | variable, to hit target $N^*$ values |
| $g_0$ (baseline growth) | 0.025 |
| $m_{\min}$ | 0.01 |
| $\alpha$ | 0.0015 |
| $\gamma$ | 0 |
| $p$ | 1.0 |
| $q$ | 0.0 |
| $m_B$ | 0.5 |
| $g_B$ | 0.5 |
| $m_0$ | 0.01–0.5 |
| *Fig. S4B, F (Case 2: growth-suppression channel, varying $m_0$)* | |
| $N^*$ values | 1, 2, 5, 10, 15, 20, 25 |
| $f$ | variable, to hit target $N^*$ values |
| $g_0$ (baseline growth) | 0.25 |
| $m_{\min}$ | 0.01 |
| $\alpha$ | 0 |
| $\gamma$ | 0.3 |
| $p$ | 1.0 |
| $q$ | 0.0 |
| $m_B$ | 0.5 |
| $g_B$ | 0.5 |
| $m_0$ | 0.01–0.2 |
| *Fig. S4C, G (Case 3: large individuals drive DD mortality* | |
| $N^*$ values | 1, 2, 5, 10, 15, 20, 25 |
| $f$ | variable, to hit target $N^*$ values |
| $m_0$ | 0.3 |
| $m_{\text{scale}}$ | 0.1 |
| $m_{\min}$ | 0.01 |
| $\alpha$ | 0.005 |
| $\gamma$ | 0 |

*(continued)*

| Parameter | Value(s) |
|---|---|
| $p$ | 1.0 |
| $q$ | 0.0 |
| $m_B$ | 0.5 |
| $g_B$ | 0.5 |
| $g_0$ | 0.01–0.5 |
| *Fig. S4D, H (Case 4: small individuals generate and experience DD mortality* | |
| $N^*$ values | 1, 2, 5, 10, 15, 20, 25 |
| $f$ | variable, to hit target $N^*$ values |
| $m_0$ | 0.3 |
| $m_{\text{scale}}$ | 0.1 |
| $m_{\text{min}}$ | 0.03 |
| $\alpha$ | 0.0001 |
| $\gamma$ | 0 |
| $p$ | $-0.8$ |
| $q$ | $-0.8$ |
| $m_B$ | 0.5 |
| $g_B$ | 0.5 |
| $g_0$ | 0.001–0.05 |

### 13 Simulation parameters for Figures 4 and 5

We provide tables of all key parameters used in the simulations presented in figures 4 and 5.

Table S2: Simulation parameters for Fig. 4 (environment-dependent measured CNDD). The environmental effect on the focal vital rate $\theta$ is implemented as $\theta(E_j) = \theta_{\text{base}} \exp(\eta_\theta E_j)$, where $E_j$ is the patch environment (standardized Gaussian field) and $\eta_\theta$ is the environmental effect size. Parameters shared across all four simulations: $n = 100$ sites, $S = 30$, $T_{\max} = 350$, $\Delta t = 5$, $b = 20$ bins, $x_{\max} = 1$, $f = 30$ (fecundity), $m_B = 0.1$ (seed mortality), $g_B = 0.2$ (germination), $\sigma_{\text{kern}} = 3$, $r_{\text{kern}} = 12$, adult autocorr.= 0.6, occupancy= 0.25, , locations= 1000.

| Parameter | Fig. 4A–B | Fig. 4C–D | Fig. 4E–F | Fig. 4G–H |
|---|---|---|---|---|
| *Simulation design* | | | | |
| Env. vital rate | $m_0$ | $m_0$ | $g_0$ | $g_0$ |
| $\eta_\theta$ | −0.6 | −0.4 | +0.5 | +0.5 |
| DD affects | Mortality ($\alpha > 0$) | Growth ($\gamma > 0$) | Mortality ($\alpha > 0$) | Mortality ($\alpha > 0$) |
| $p$ | 1.0 | 1.0 | 1.0 | −0.8 |
| $q$ | 0.0 | 0.0 | 0.0 | −0.8 |
| *Baseline species parameters* | | | | |
| $m_0$ | 0.1 | 0.1 | 0.1 | 0.1 |
| $m_{\min}$ | 0.01 | 0.01 | 0.01 | 0.03 |
| $g_0$ | 0.06 | 0.25 | 0.06 | 0.06 |
| $\alpha_{\text{con}}$ | 0.002 | 0.0 | 0.002 | 0.0001 |
| $\alpha_{\text{het}}$ | 0.0004 | 0.0 | 0.0004 | 0.00002 |
| $\gamma_{\text{con}}$ | 0.0 | 1.0 | 0.0 | 0.0 |
| $\gamma_{\text{het}}$ | 0.0 | 0.2 | 0.0 | 0.0 |
| *Interspecific variation (lognormal CV)* | | | | |
| $\text{CV}_{m_0}$ | 0.3 | 0.3 | 0.3 | 0.3 |
| $\text{CV}_{g_0}$ | 0.3 | 0.3 | 0.3 | 0.3 |
| $\text{CV}_\alpha$ | 0.3 | — | 0.3 | 0.3 |
| $\text{CV}_\gamma$ | — | 0.3 | — | — |

Table S3: Simulation parameters for Fig. 5. *Top*: four main scenarios (panels A–H). *Bottom*: variance-decomposition conditions (panel I), all using M-channel ($\alpha > 0$, $\gamma = 0$), $p = 1$, $q = 0$, $k = 1$. Conditions 2–4 copy each species' full parameter vector from "Both vary", overriding only the fixed vital rate(s) to their baseline value. Parameters shared across all simulations: $n = 100$ sites, $S = 75$, $T_{\max} = 350$, $\Delta t = 5$, $b = 20$ bins, $x_{\max} = 1$, $f = 30$ (fecundity, except Fig. 5C–D where $f = 10$), $m_B = 0.1$ (seed mortality), $g_B = 0.2$ (germination), $\sigma_{\text{kern}} = 3$, $r_{\text{kern}} = 12$, adult autocorr.= 0.6, occupancy= 0.25, $m_{\min} = 0.01$, locations= 1000.

| | *Main scenarios (A–H)* | | | | *Variance decomposition (I)* | | | |
|---|---|---|---|---|---|---|---|---|
| Parameter | 5A–B | 5C–D | 5E–F | 5G–H | Both vary | Only $m_0$ | Only $g_0$ | Neither |
| DD affects | Mortality ($\alpha > 0$) | Growth ($\gamma >$ 0) | Mortality ($\alpha > 0$) | Mortality ($\alpha > 0$) | Mortality ($\alpha > 0$) | Mortality ($\alpha > 0$) | Mortality ($\alpha > 0$) | Mortality ($\alpha > 0$) |
| $p$ | 1.0 | 1.0 | 1.0 | −0.9 | 1.0 | 1.0 | 1.0 | 1.0 |
| $q$ | 0.0 | 0.0 | 0.0 | −0.9 | 0.0 | 0.0 | 0.0 | 0.0 |
| $m_0$ | 0.1 | 0.1 | 0.1 | 0.1 | 0.1 | 0.1 | 0.1 | 0.1 |
| $g_0$ | 0.06 | 0.25 | 0.06 | 0.06 | 0.06 | 0.06 | 0.06 | 0.06 |
| $\alpha_{\text{con}}$ | 0.02 | 0.0 | 0.02 | 0.0001 | 0.01 | 0.01 | 0.01 | 0.01 |
| $\alpha_{\text{het}}$ | 0.004 | 0.0 | 0.004 | 0.00002 | 0.002 | 0.002 | 0.002 | 0.002 |
| $\gamma_{\text{con}}$ | 0.0 | 1.0 | 0.0 | 0.0 | 0.0 | 0.0 | 0.0 | 0.0 |
| $\gamma_{\text{het}}$ | 0.0 | 0.2 | 0.0 | 0.0 | 0.0 | 0.0 | 0.0 | 0.0 |
| Fecundity | 30 | 10 | 30 | 30 | 30 | 30 | 30 | 30 |
| Focal param. | $m_0$ | $m_0$ | $g_0$ | $g_0$ | $m_0$, $g_0$ | $m_0$ | $g_0$ | — |
| $\text{CV}_{m_0}$ | $0.75^{\dagger}$ | $0.60^{\dagger}$ | 0.20 | 0.20 | $0.75^{\dagger}$ | $0.75^{\dagger}$ | fixed | fixed |
| $\text{CV}_{g_0}$ | 0.20 | 0.10 | $0.75^{\dagger}$ | $0.75^{\dagger}$ | $0.75^{\dagger}$ | fixed | $0.75^{\dagger}$ | fixed |
| $\text{CV}_{\alpha}$ | 0.20 | — | 0.20 | 0.20 | — | — | — | — |
| $\text{CV}_{\gamma}$ | — | 0.05 | — | — | — | — | — | — |

$^{\dagger}$ Focal parameter (varied at high CV to drive interspecific variation in CNDD).

### 14 Variable / parameter table

For reference, we provide a single table of all symbols (parameters, variables, etc.) used across the main text and supplement.

Table S4: Notation and parameters used in the main text and Supplement.

| Symbol / term | Meaning | Notes / domain |
|---|---|---|
| *General notation* | | |
| $t$ | Continuous time. | $t \geq 0$ |
| $T$ | Census interval length. | $T > 0$ |
| $x$ | Individual size. | $x \in [0, x_{\max}]$ |
| $x_{\max}$ | Maximum size. | |
| $N(0)$ | Initial cohort size (interpreted as local conspecific density). | $N(0) > 0$ |
| $N(t)$ | Total cohort abundance at time $t$. | |
| $N^*$ | Equilibrium (steady-state) cohort abundance. | |
| $p_i$ | Survival probability of individual/replicate $i$ over $[0, T]$; $p_i = \exp(-\mathrm{H}(T))$. | |
| $\mathrm{H}(T)$ | Cohort cumulative hazard over $[0, T]$; $\mathrm{H}(T) = -\log(N(T)/N(0))$. | |
| $\bar{h}(t)$ | Abundance-weighted mean instantaneous hazard; written $\bar{m}(t)$ when referring specifically to mortality in the SI and PDE models. | |
| $\mathrm{H}_{\mathrm{DD}}(T)$ | Cumulative excess hazard attributable to density dependence; $\mathrm{H}_{\mathrm{DD}}(T) = \mathrm{H}(T) - \mathrm{H}_{\mathrm{DI}}(T)$. This is realized interval CNDD. | |
| $\mathrm{H}_{\mathrm{DI}}(T)$ | Cumulative density-independent hazard over $[0, T]$, from a matched simulation with density dependence removed ($\beta = 0$ or $\alpha = \gamma = 0$). | |
| $\bar{m}_{\mathrm{DI}}(t)$ | Cohort-averaged density-independent hazard at time $t$ (density dependence removed). | |
| $\beta_1$ | Generic logit-scale slope of survival on density (negative = CNDD); instantiated as $\beta_1(m)$ in the SI model or $\beta_{\mathrm{con}}$ in the GLMM. | $\leq 0$ |
| *Key conceptual quantities* | | |
| $\alpha$ | Instantaneous density sensitivity of mortality. | $\alpha \geq 0$ |
| $\gamma$ | Instantaneous density sensitivity of growth. | $\gamma \geq 0$ |
| Realized interval CNDD | $\mathrm{H}_{\mathrm{DD}}(T)$, the cumulative survival consequence of crowding over $[0, T]$; its density-sensitivity $\partial \mathrm{H}_{\mathrm{DD}}(T)/\partial N(0)$ is what drives $\beta_1$. | |
| Measured CNDD | Statistical coefficient estimated from interval-census data ($\hat{\beta}_1(m)$, $\beta_{\mathrm{con}}$, or $u_{\mathrm{con},s}$). | |
| *SI pathogen model — cohort dynamics* | | |
| $S(t)$ | Number of susceptible individuals at time $t$. | integer $\geq 0$ |
| $I(t)$ | Number of infected individuals at time $t$. | integer $\geq 0$ |
| $\xi$ | Background infection rate (exposure independent of $I$). | $\xi > 0$ |

*(continued)*

| Symbol / term | Meaning | Notes / domain |
|---|---|---|
| $\beta$ | Mass-action transmission rate. | $\beta > 0$; distinct from regression $\beta_1$ (see Methods note) |
| $m$ | Baseline (density-independent) per-capita mortality rate. | $m > 0$ |
| $m_{\text{inf}}$ | Additional per-capita mortality rate of infected individuals. | $m_{\text{inf}} > 0$ |
| $f$ | Seedling supply rate (SI model) / seed input rate (PDE model). | shared symbol across both models |
| $\lambda_{\text{infection}}$, $\lambda_{\text{death},S}$, $\lambda_{\text{death},I}$ | Gillespie SSA event hazards for infection and death events. | |
| $y_k$ | Survival outcome of SSA replicate $k$. | $\in \{0, 1\}$ |
| $p_k$ | Modeled survival probability of replicate $k$ in the SI GLM. | |
| $\beta_0(m)$ | GLM intercept, as a function of $m$. | |
| $\hat{\beta}_1(m)$ | Fitted GLM slope of survival on $N(0)$; measured CNDD in the SI model. | $\leq 0$ |
| *SI pathogen model — establishment probability* | | |
| $R_0$ | Basic reproduction number; $R_0 = \beta N(0)/(m + m_{\text{inf}})$. | threshold at $R_0 = 1$ |
| $R_t$ | Effective reproduction number at time $t$; $R_t = R_0 e^{-mt}$. | decreasing in $t$ |
| $t_c$ | Critical time beyond which $R_t < 1$; $t_c = (1/m)\log R_0$. | |
| $\lambda_{\text{intro}}(t)$ | Rate of background introductions; $\lambda_{\text{intro}}(t) = \xi S(t)$. | |
| $q(t)$ | Establishment probability given introduction at time $t$. | $\in [0, 1]$; note $q$ is reused as the size-susceptibility exponent below (unrelated) |
| $r(t)$ | Intensity of the thinned Poisson process of successful establishments; $r(t) = \lambda_{\text{intro}}(t)\, q(t)$. | |
| $\tau$ | Effective upper limit of integration; $\tau = \min(T, t_c)$. | |
| $P_{\text{est}}(T)$ | Probability of infection establishment by census time $T$. | $\in [0, 1]$ |
| *SI pathogen model — equilibrium (supply) variant* | | |
| $S^*$, $I^*$ | Endemic equilibrium susceptible / infected abundances. | |
| $m^*_{\text{DD}}$ | Equilibrium excess per-capita mortality hazard due to infection; $m^*_{\text{DD}} = m_{\text{inf}} I^*/N^*$. | |
| $T_{\text{burn}}$ | Burn-in duration used to reach steady state before tracking a census interval. | |
| *Size-structured seedling model* | | |
| $n(x, t)$ | Seedling size density; $n(x, t)\, dx$ is the number of individuals with size in $[x, x + dx]$. | |
| $B(t)$ | Local seed pool abundance at time $t$. | |
| $m_B$ | Per-capita seed loss rate. | $m_B > 0$ |
| $g_B$ | Per-capita germination rate. | $g_B > 0$ |
| $g(x, t)$ | Per-capita growth rate of an individual of size $x$ at time $t$. | |
| $m(x, t)$ | Per-capita mortality hazard of an individual of size $x$ at time $t$. | |
| $g_0$ | Baseline growth rate parameter. | $g_0 > 0$ |
| $m_0$ | Mortality hazard at smallest size ($x = 0$). | $m_0 > 0$ |

*(continued)*

| Symbol / term | Meaning | Notes / domain |
|---|---|---|
| $m_{\min}$ | Mortality hazard at largest size ($x = x_{\max}$). | $m_0 > m_{\min} \geq 0$ |
| $m_{\text{scale}}$ | Scalar multiplier on overall mortality level (used only in Supp. parameter tables; not formally introduced by an equation — flagged). | |
| $m_{\text{DI}}(x)$, $g_{\text{DI}}(x)$ | Density-independent components of mortality/growth (Eqs. MortEq/GrowthEq). | |
| $m_{\text{DD}}(x,t)$, $g_{\text{DD}}(x,t)$ | Density-dependent components of mortality/growth. | |
| $P(t)$ | Neighborhood crowding index; $P(t) = \int_0^{x_{\max}} w(x)\, n(x,t)\, dx$. | |
| $w(x)$ | Size-dependent neighbor weighting function; $w(x) = x^p$. | |
| $p$ | Neighbor weighting exponent; $p > 0$ upweights larger neighbors, $p < 0$ upweights smaller. | |
| $\phi_M(x)$, $\phi_G(x)$ | Size-dependent susceptibility functions for the mortality/growth channels; $\phi(x) = (x/x_{\max})^q$. | |
| $q$ | Susceptibility exponent; $q < 0$ implies stronger effects on smaller individuals. | |
| $P^*$ | Equilibrium (standing) crowding field. | |
| $n^*(x)$ | Equilibrium size distribution. | |
| *Two-stage approximation (Box 1)* | | |
| $J_1(t)$, $J_2(t)$ | Abundances of the "small" and "large" stages. | $J_1(0) = N(0)$, $J_2(0) = 0$ |
| $g(J_1, J_2)$ | Per-capita transition rate from small to large stage; baseline value $g_0$ (Cases 1,3,4) or $g_0 \exp[-\gamma(J_1 + J_2)]$ (Case 2). | |
| $\mu_1(J_1, J_2)$, $\mu_2(J_1, J_2)$ | Per-capita mortality hazards of the small/large stage. | |
| $m_1$, $m_2$ | Baseline (density-independent) mortality hazards of the small/large stage. | $m_1 > m_2$ typical |
| *Spatially explicit multispecies simulations* | | |
| $s$ | Species index. | |
| $j$ | Patch index. | |
| $E_j$ | Environmental value of patch $j$. | standardized |
| $\theta$ | Generic vital-rate parameter subject to environmental variation ($m_0$ or $g_0$). | |
| $\theta_{\text{baseline}}$ | Baseline (reference) value of $\theta$. | |
| $\eta_\theta$ | Environmental effect size on $\theta$; $\theta(E_j) = \theta_{\text{baseline}} \exp(\eta_\theta E_j)$. | |
| $n$ | Grid side length; total patches = $n^2$. | |
| $S$ | Total number of species in a simulation. | |
| $T_{\max}$ | Total simulated time horizon. | |
| $\Delta t$ | Length of each recorded census interval. | |
| $b$ | Number of size bins in the discretization. | integer |
| $\Delta x$ | Bin width; $\Delta x = x_{\max}/b$. | |
| $\ell$ | Size-bin index; $x_\ell = (\ell - \frac{1}{2})\Delta x$, $\ell = 1, \ldots, b$. | |
| $N_{s,\ell}(t)$ | Integer abundance of species $s$ in size bin $\ell$ at time $t$. | |
| $P_{\text{con},s}(t)$, $P_{\text{het},s}(t)$ | Conspecific / heterospecific crowding index for species $s$. | |
| $\alpha_{\text{con},s}$, $\alpha_{\text{het},s}$ | Conspecific / heterospecific instantaneous density sensitivity of mortality for species $s$. | |

*(continued)*

| Symbol / term | Meaning | Notes / domain |
|---|---|---|
| $\gamma_{\text{con},s}$, $\gamma_{\text{het},s}$ | Conspecific / heterospecific instantaneous density sensitivity of growth for species $s$. | |
| $\sigma_{\text{kern}}$ | Dispersal kernel standard deviation. | appears only in Table S4/S5 captions; please confirm description |
| $r_{\text{kern}}$ | Dispersal kernel radius | Truncates seed dispersal (for numerical ease) |
| $\text{CV}_\theta$ | Coefficient of variation of the lognormal interspecific distribution for vital rate $\theta$. | Table S5 |
| *Statistical models (GLM/GLMM) for simulated censuses* | | |
| $y_i$ | Survival outcome of individual $i$ in a census window. | $\in \{0, 1\}$ |
| $\beta_0$ | GLMM intercept. | |
| $\text{Size}_i$ | Initial size of individual $i$. | |
| $\beta_{\text{size}}$ | Fixed effect of initial size. | |
| $\text{Con}_i$, $\text{Het}_i$ | Power-transformed, z-scored conspecific / heterospecific crowding covariates. | |
| $\text{Con}_{i,\text{raw}}$, $\text{Het}_{i,\text{raw}}$ | Raw (untransformed) conspecific / heterospecific crowding counts. | |
| $z(\cdot)$ | Z-scoring (standardization to mean 0, unit variance) within the fitted dataset. | |
| $b_1$, $b_2$ | Power-transform exponents applied to $\text{Con}_{i,\text{raw}}$, $\text{Het}_{i,\text{raw}}$. | grid-searched over $\{0.25, \ldots, 2\}$ |
| $\beta_{\text{con}}$, $\beta_{\text{het}}$ | Population-level fixed effects of conspecific / heterospecific density on survival (logit scale). | $\leq 0$ for CNDD |
| $u_{0,s}$ | Species-level random intercept. | |
| $u_{\text{con},s}$, $u_{\text{het},s}$ | Species-level random slopes for conspecific / heterospecific density; species-level measured CNDD. | |
| $\widehat{\beta}_{\text{con},s}$ | Species-level conspecific density slope, $\beta_{\text{con}} + u_{\text{con},s}$; the total measured CNDD for species $s$. | |
| $E_{j(i)}$ | Environment of the patch containing individual $i$. | |
| $\beta_E$ | Fixed effect of patch environment. | |
| $\beta_{E:\text{con}}$, $\beta_{E:\text{het}}$ | Environment-by-conspecific / environment-by-heterospecific interaction coefficients. | |
| $K$ | Number of final census intervals retained for analysis (transient exclusion). | $K = 20$ in Figs. 4–5 |
